# The glycolysis regulator PFKFB4 interacts with ICMT and activates RAS/AKT signaling-dependent cell migration in melanoma

**DOI:** 10.1101/2020.03.23.004119

**Authors:** Méghane Sittewelle, Vincent Kappès, Déborah Lécuyer, Anne H. Monsoro-Burq

## Abstract

Cell migration is a complex process, tightly regulated during embryonic development and abnormally activated during cancer metastasis. RAS-dependent signaling is a major nexus controlling essential cell parameters including proliferation, survival and migration, utilizing downstream effectors such as the PI3K/AKT signaling pathway. In melanoma, oncogenic mutations frequently enhance RAS, PI3K/AKT or MAP kinase signaling, and trigger other cancer hallmarks among which the activation of metabolism regulators. PFKFB4 is one of these critical regulators of glycolysis and of the Warburg effect. Here however, we explore a novel function of PFKFB4 in melanoma cell migration. We find that PFKFB4 interacts with ICMT, a post-translational modifier of RAS. PFKFB4 promotes ICMT/RAS interaction, controls RAS localization at the plasma membrane, activates AKT signaling and enhances cell migration. We thus provide evidence of a novel and glycolysis-independent function of PFKFB4 in human cancer cells. This unconventional activity links the metabolic regulator PFKFB4 to RAS-AKT signaling and impacts melanoma cell migration.

**Highlights:**

- PFKFB4, a known regulator of glycolysis, also displays an unconventional role in melanoma cell migration.
- PFKFB4 interacts with ICMT and promotes RAS localization at the plasma membrane.
- PFKFB4 and ICMT cooperation modulates AKT signaling and controls melanoma cell migration.

## Introduction

Cell migration is one of the critical processes involved in the formation and maintenance of multicellular organisms. To acquire motility, cells activate complex properties such as cytoskeleton remodeling, inhibition of cell-cell contacts, remodeling the extracellular matrix and response to chemo-attractants (*1, 2*). To orchestrate these processes, numerous redundant and complementary signaling pathways cooperate with one another. While many of these pathways have been studied separately, the crosstalk between signaling and other parameters, such as cell cycle or cell metabolism, only begins to be explored in stem cells, normal development and cancer (*3*). Cell motility and invasiveness properties are reactivated during cancer progression, following the aberrant activation of multiple cellular programs, such as growth factor-independent signaling, metabolic and epigenetic reprograming, which cooperate to sustain growth, proliferation, and survival properties in the primary tumor (*3*). Cancer cell migration, metastasis and formation of secondary tumors are the major cause of death for aggressive cancers such as cutaneous melanoma, the deadliest skin cancer in humans (*4*). The mechanisms driving melanoma cell invasion are multiple and remain incompletely understood. To treat melanoma, one possible therapeutic strategy is to target identified driver mutations. The numerous genetic alterations involved in cutaneous melanoma development can be classified into four subtypes: BRAF, RAS (N/H/K), NF1 and Triple-WT (*5*). The mitogen-activated protein kinase (MAPK) pathway is the most frequently altered with approximately 50% of tumors mutated in *BRAF* gene, followed by 25% with mutations in NRAS and 14% in NF1, leaving 10% of the tumors with no identified driver mutation and more complex genetic landscapes (*5, 6*). Although RAS signaling acts upstream of both MAPK and PI3K/AKT signaling (*7*), it is interesting to note that the two main *NRAS* activating mutations in cutaneous melanoma, the hotspots Q61 (80% of *NRAS* subtype tumors) and G12 (15%), drive a differential activation of the downstream pathways with preferential activation of MAPK or PI3K/AKT respectively, suggesting a complex modulation of the structure and activity of oncogenic proteins (*8*). In all cases, these alterations in signaling lead to increased melanoma cell proliferation, survival and migration.

In addition to mutations activating signaling pathways, metabolism rewiring allows cancer cells to promote active cell proliferation. In particular, enhanced glycolysis rate even under normal oxygen conditions, called the Warburg effect, drives many parallel biosynthesis pathways to provide cellular building blocks together with bioenergy (*9*). Moreover, the hypoxic environment often found in early primary tumors prior to vascularization, also stimulates the activation of metabolic regulators induced by the hypoxia-inducing factor HIF1. This is the case for the family of 6-phosphofructo-2-kinase/fructose-2,6-biphosphatases enzymes (PFKFB1-4), which are major regulators of glycolysis, controlling the rate of the second irreversible and rate-limiting reaction of glycolysis catalyzed by the phospho-fructokinase 1 (PFK1) (*10*). PFKFB enzymes are bi-functional and synthesize (with kinase activity) or degrade (with phosphatase activity) the fructose-2,6-biphosphate, the main allosteric activator of PFK1. Thus, increased PFKFB protein kinase activity promotes glycolysis. In human, four distinct genes encode PFKFB isoenzymes 1 to 4, each one possessing many splicing isoforms and differing in their tissue-specific abundance, kinetics and regulation properties (*11–16*). PFKFB proteins are overexpressed in cancer. In particular, increased PFKFB4 levels have been reported in several human tumors, including cutaneous melanoma (*17–20*). Moreover, *PFKFB4* is induced by hypoxia, is required for survival and proliferation of normal thymocytes (*21*), as well as of several cancer cell lines such as lung, breast and colon adenocarcinomas, prostate and bladder cancer (*22–25*). So far, most studies have focused on cell metabolism reprogramming by PFKFB4 and have proposed that PFKFB4 is a major driver of Warburg effect (*17, 19, 20, 22–30*). However, a few recent studies have identified alternative functions of PFKFB4, outside of its canonical control of glycolysis. For example, PFKFB4 regulates small lung-cancer chemo-resistance by stimulating autophagy, via its interactions with Etk tyrosine kinase (*31, 32*). PFKFB4 also operates as a protein kinase and directly phosphorylates SRC-3, promoting metastatic progression in highly glycolytic breast cancer cells (*33*). During development, PFKFB4 is essential for early embryonic inductions and neural crest cells migration through the activation of AKT signaling (*34, 35*). In cancer, the intriguing relationships between PFKFB4, cell signaling and cell migration remain unexplored.

Here we have analyzed the importance of PFKFB4 in melanoma cell migration. Using human metastatic melanoma cell lines with high *PFKFB4* expression (*36*), we show that PFKFB4 activity is required for active cell migration in several different cellular contexts, without a connection to the rate of glycolysis. Rather, we identify potential interacting proteins by mass spectrometry, among which we validate the protein-protein interactions between PFKFB4 and isoprenylcystein carboxymethyl transferase (ICMT), an enzyme essential for RAS post-translational modifications controlling its localization at the plasma membrane. Our study further defines a novel, glycolysis-independent function for PFKFB4, which promotes ICMT-RAS interactions, results in efficient RAS localization at the plasma membrane, activates AKT signaling and enhances melanoma cell migration.

## Results

### PFKFB4 controls metastatic melanoma cell migration in vitro in a glycolysis-independent manner

Melanomas present higher expression of *PFKFB4* compared to other tumors (Figure S1). We have previously linked elevated expression of PFKFB4 with embryonic cell migration *in vivo* (*35*), but in melanoma, while PFKFB4 has been linked to promoting the Warburg effect, its role in cell migration remains to be explored. Here we have chosen two human melanoma cell lines expressing high levels of PFKFB4 (MeWo and A375M, Figure 1A-B, (*36*)) to follow the random migration of individual cells by time-lapse video microscopy (Figure 1D). The MeWo cells are derived from lymph node metastasis of a cutaneous melanoma. They are tumorigenic and metastatic. They bear wild-type alleles at BRafV600 or RasQ61/G12 positions (*36, 37*) (Figure S2A). The A375M cells are derived from a human amelanotic melanoma. They are also tumorigenic and metastatic. They are mutated for BRafV600 and wild-type for RasQ61 (*36*). Both cell lines actively migrated on Matrigel. Cells were tracked during 16 hours in at least three independent experiments for each cell line. After PFKFB4 depletion using siRNA, both MeWo and A375M cells migrated in average twice slower than control (Figure 1E,F; Table S1). Migration distance was also decreased while cell pausing was increased (Figure S2C-F). Because during *X. laevis* embryonic development, the migration of melanocytes and melanoma progenitors, the neural crest cells, is controlled by PFKFB4 (*35*), we postulated that human and frog protein functions were conserved, allowing us to devise phenotype rescue experiments: Frog *pfkfb4* encodes a protein with 95% similarity with the human protein, but the mRNA was not targeted by siRNAs designed against the human mRNA sequence. In MeWo cells, the migration phenotype was efficiently rescued by co-transfecting the *Xenopus laevis* orthologous *pfkfb4* sequence (Figures 1E-F, S2C). This rescue showed that the migration phenotype was specific for the depletion of PFKFB4 protein and did not come from off-target effects.

**Figure 1.**
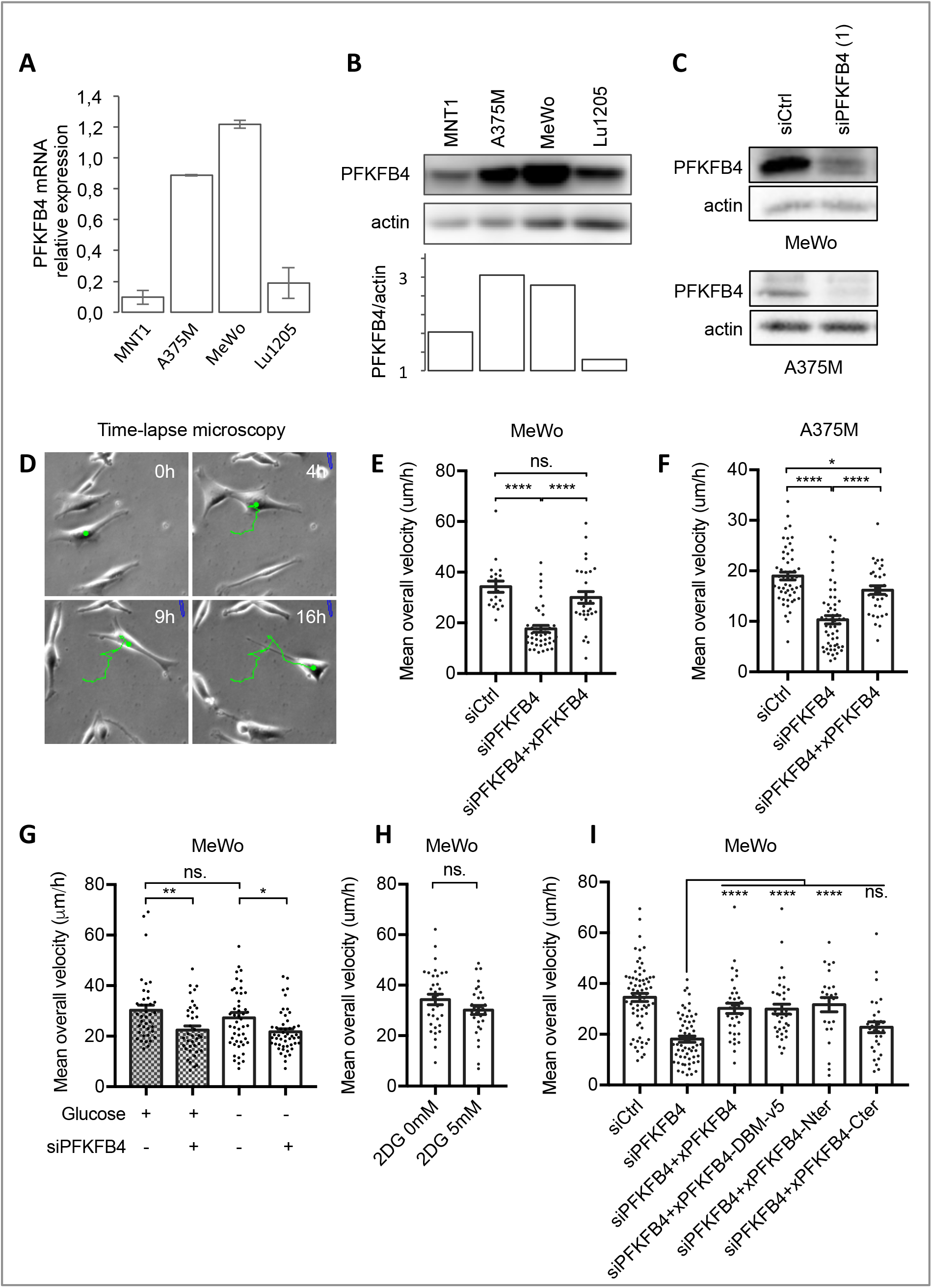
PFKFB4 controls in vitro cell migration in metastatic melanoma in a glycolysis-independent manner. (A,B) Quantification of PFKFB4 mRNA and protein levels in four human melanoma cell lines: MNT1, A375M, MeWo and Lu1205. (A) Relative *PFKFB4* mRNA levels measured by RT-qPCR and normalized by the expression of 18S and TBP housekeeping genes. Error bars: S.E.M. (B) PFKFB4 protein levels detected by Western-blotting, with PFKFB4/actin relative quantification (optical density). (C) PFKFB4 protein levels in MeWo and A375M cells 48h after transfection with siRNA targeting PFKFB4. (D) Starting 48h after transfection, cell migration was tracked for 16 hours from phase contrast images. Each point corresponds to the average speed of one cell. MeWo (E,G,H,I) or A375M (F) cells were co-transfected either with siControl/empty plasmid, siPFKFB4/empty plasmid or with siPFKFB4 together with a *X. laevis* PFKFB4 plasmid in its wild-type form (E,F,I) or mutant forms (I). The average speed was also measured when cells were cultured in glucose-free medium (G) or complete medium supplemented with 2DG (H). In each panel, a representative experiment is shown (n>3), and displays mean ± SEM. p-values were calculated using the Mann-Whitney test. n.s.: p > 0,05; *: p < 0,05; **: p < 0.01; ***: p < 0.001; ****: p < 0.0001.

We then investigated if PFKFB4’s role in cell migration was linked with its function as a major activator of the glycolysis rate. To block glycolysis, we first grew the MeWo cells in a culture medium without glucose, which strongly decreased their glycolysis rate (estimated by the diminished lactate production measured in the culture medium, Figure S3B). We did not observe a major decrease of MeWo cells’ average migration speed in the glucose-free medium compared to the complete medium condition (Figure 1G,H). Next, to confirm that MeWo cell migration was unaffected by limiting glycolysis, we added the glucose non-hydrolysable analog 2-deoxyglucose (2DG) in the complete medium (Figure S3A). Similar to the glucose-free condition, 2DG also decreased glycolysis efficiently (Figure S3C) without affecting MeWo cells’ migration speed (5 mM 2DG, Figure 1H). These results indicated that MeWo cells’ migration was not directly linked to their rate of glycolysis. In contrast, PFKFB4-depleted MeWo cells showed a reduced average migration speed compared to control cells in the glucose-free medium as observed in the complete medium, indicating an action of PFKFB4 on another cellular pathway (Figure 1G). Together, these observations suggested that PFKFB4 levels significantly affect the average speed of cells migration, independently of the rate of glycolysis.

These observations were in contrast with the expected phenotype, as the best-known function of the bi-functional enzyme PFKFB4 is to regulate the glycolysis rate, by controlling the second rate-limiting reaction of glycolysis. With its kinase moiety, PFKFB4 phosphorylates fructose-6-phosphate into fructose-2,6-bisphosphate, the allosteric activator of PFK1 (*14, 38*) (Figure S3A). With its phosphatase domain, PFKFB4 catalyzes the reverse reaction. To understand if PFKFB4 controlled melanoma cells migration using either its kinase or its phosphatase enzymatic activities, we compared the rescue phenotype of PFKFB4 depletion by various *X. laevis* PFKFB4 mutants (Figure S2B). The average speed of cells co-transfected with siPFKFB4 and a plasmid encoding a *pfkfb4* mutant with two point mutations inactivating both kinase and phosphatase enzymatic activities simultaneously (mutations G48A and H258A, called xPFKFB4-DBM; (*39, 40*)) was equivalent to the speed after rescue by wild-type PFKFB4 (xPFKFB4) (Figure 1I). This indicated that PFKFB4 controlled cell migration independently of its enzymatic activities. To identify which region of PFKFB4 protein was involved in this non-conventional effect, we used two complementary deletion constructs. The rescue done with a deletion construct encoding the N-terminal kinase domain (xPFKFB4-Nter) was as efficient as with xPFKFB4 (Figure 1I). In contrast, the migration of cells depleted for PFKFB4 and co-transfected with the deletion construct encoding only the C-terminal phosphatase domain (xPFKFB4-Cter) was not rescued (Figure 1I). These results suggested that PFKFB4 was involved in control of cell migration independently of its kinase or phosphatase activities, but through the N-terminal half of the protein.

### PFKFB4 interacts with ICMT, a major post-translational modifier of RAS GTPases

As PFKFB4 seemed to control melanoma cell migration independently of its enzymatic activities, we looked for interacting protein partners using mass spectrometry after immunoprecipitation of a FLAG-tagged form of human PFKFB4 expressed in the MeWo cells. In two biological replicates, among 1556 high confidence hits, we chose 41 candidates with a Mascot score enriched at least ten-times compared to the negative control condition in order to eliminate the weak hits and limit the non-specific targets. Moreover, because xPFKFB4 efficiently rescued the PFKFB4 depletion phenotype in human melanoma cells, we postulated that the protein function we looked for was evolutionarily conserved between human and *X. laevis* PFKFB4. We transfected MeWo cells with the frog xPFKFB4 ortholog followed by immunoprecipitation and mass spectrometry. We then crossed the 41-candidates list obtained with hPFKFB4 with the list of xPFKFB4 targets and sub-selected 23 candidates (Figures 2A, S4). Among these 23 best candidates, we prioritized isoprenylcystein carboxyl methyl transferase (ICMT), a potential modulator of PI3K/AKT signaling pathway, because PFKFB4 was known to affect cell migration via AKT signaling activation during embryogenesis (*34, 35*). ICMT is an endoplasmic reticulum membrane protein critical for RAS GTPases post-translational modifications. ICMT catalyzes the carboxyl methylation of RAS on its C-terminal CAAX motif. This modification allows RAS protein to be targeted to the plasma membrane, a prerequisite for the coordination by RAS of a variety of signaling pathways, including PI3K/AKT activation ((*41–44*), reviewed in (*45*)). This observation suggested that an interaction between PFKFB4 and ICMT could occur during melanomagenesis and be related to RAS-dependent signaling pathways.

**Figure 2.**
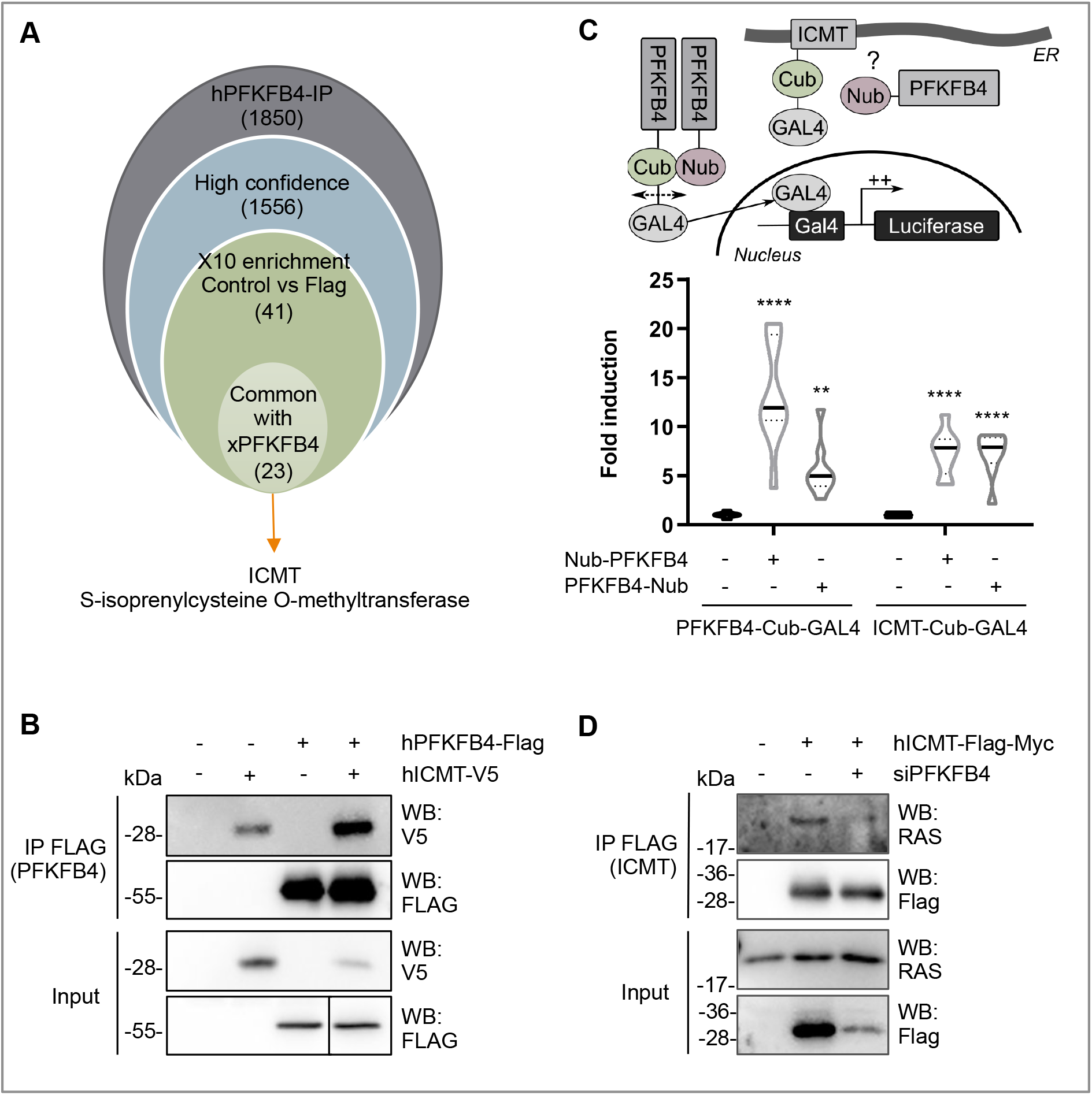
PFKFB4 interacts with ICMT, a major post-translational modifier of RAS GTPases. (A) Workflow used to select candidates after PFKFB4 immunoprecipitation followed by mass spectrometry analysis (see text for details). (B) Enrichment of ICMT tagged with V5 after immunoprecipitation by FLAG-PFKFB4 from MeWo cell extracts. (C) Scheme of the MaMTH strategy used to validate PFKFB4/ICMT protein-protein interactions. Violin plot showing the luciferase activity measured and normalized from MaMTH modified HEK293T cells extracts (n=3). The violin represents the probability density at each value, lines are plotted at the median and quartiles (Two-way ANOVA test. **: p < 0.01 and ****: p < 0.0001.). (D) The interaction between tagged ICMT and endogenous RAS was evaluated with or without PFKFB4 depletion. Immunoprecipitation of FLAG-ICMT from MeWo cells followed by Western-blotting with antibody against V5, FLAG or endogenous RAS.

First, we validated the mass spectrometry results by co-immunoprecipitation of ICMT with PFKFB4 in MeWo cells (Figure 2B). Secondly, we confirmed the protein-protein interaction between PFKFB4 and ICMT by an independent alternative method, a split-ubiquitin two-hybrid approach adapted for mammalian membrane proteins (MaMTH) (Figure 2C) (*46, 47*). Briefly, ICMT and PFKFB4 were fused to a portion of the ubiquitin protein, either its N-ter part (Nub, <u>N-ub</u>iquitin) or its C-ter part fused to GAL4 (Cub-GAL4, <u>C-ub</u>iquitin). Constructs were co-transfected in MaMTH-modified HEK293T cells bearing a stable integration of GAL4-binding sites upstream of a luciferase reporter. Upon interaction, the two halves of ubiquitin reunite to form a “pseudo-ubiquitin” which recruits deubiquitinating enzymes (DUBs). The DUBs then cleave the pseudo-ubiquitin, resulting in the release of the GAL4 transcription factor. GAL4 then activates the transcription of GAL4-driven luciferase in the nucleus. As a positive control, co-transfection of PFKFB4-Cub-GAL4 and PFKFB4-Nub strongly increased luciferase expression (by five to thirteen-times compared to PFKFB4-Cub-GAL4 alone, for PFKFB4-Nub fusion in N-ter or C-ter respectively). This denoted a strong and stable interaction, related to the formation of the PFKFB4 homodimer. The co-transfection of ICMT-Cub-GAL4 and PFKFB4-Nub significantly increased luciferase expression compared to ICMT-Cub-GAL4 alone and in a range comparable to the known PFKFB4-PFKFB4 homophilic interaction. Together, these results demonstrated that PFKFB4 and ICMT directly interacted. Lastly we tested if the interaction between PFKFB4 and ICMT was important for the known interaction between ICMT and RAS GTPase. When PFKFB4 was depleted in MeWo cells, we observed a decrease of endogenous RAS immunoprecipitation by ICMT (Figure 2D). In sum, all these results suggested that PFKFB4 direct protein-protein interactions with ICMT impacted ICMT-RAS complex formation in melanoma.

### ICMT and PFKFB4 control RAS localization at the plasma membrane and melanoma cell migration

To understand the role of the PFKFB4-ICMT interaction, we first compared PFKFB4 and ICMT depletion phenotypes in MeWo cells, using a validated siRNA against ICMT (*45*) (Figure S5A). Parameters of melanoma cell migration were measured as mentioned previously. Compared to control siRNA, cells transfected with siICMT exhibited a decrease in their average speed of migration, as well as altered pausing and distance parameters, similar to cells transfected with siPFKFB4 (Figure 3A, Figure S5B,C). To test the interdependency of PFKFB4 and ICMT, we co-transfected both siRNAs. Melanoma MeWo cells receiving both siPFKFB4 and siICMT did not exhibit a more severe phenotype than with either siRNA alone. This suggested that PFKFB4 and ICMT cooperated in the same pathway to control cell migration and that depleting either one was sufficient for attaining a strong phenotype (Figure 3A). To further test this hypothesis, we tested the epistasis between PFKFB4 and ICMT by combining depletion of one factor and gain-of-function of the other, in order to see if increased activity of either one of these proteins could compensate for the loss of the other, as could be the case if they were acting in parallel and redundant pathways: the siPFKFB4 was co-transfected with the *ICMT* expression plasmid, or the siICMT with the *PFKFB4* expression plasmid. When compared to the migration speed of MeWo cells transfected with a control siRNA and the corresponding siRNA alone, neither ICMT nor PFKFB4 gain of function rescued the phenotype of PFKFB4 or ICMT depletion respectively (Figure 3B). This suggested two alternative (and not exclusive) possibilities: either the need for both proteins simultaneously, cooperating to enhance cell migration, or that one of these two proteins was functional only after being activated by the other. As a whole, this series of results showed that ICMT depletion phenocopied loss of PFKFB4, and that the two protein partners were likely acting in the same pathway impacting melanoma cell migration.

**Figure 3.**
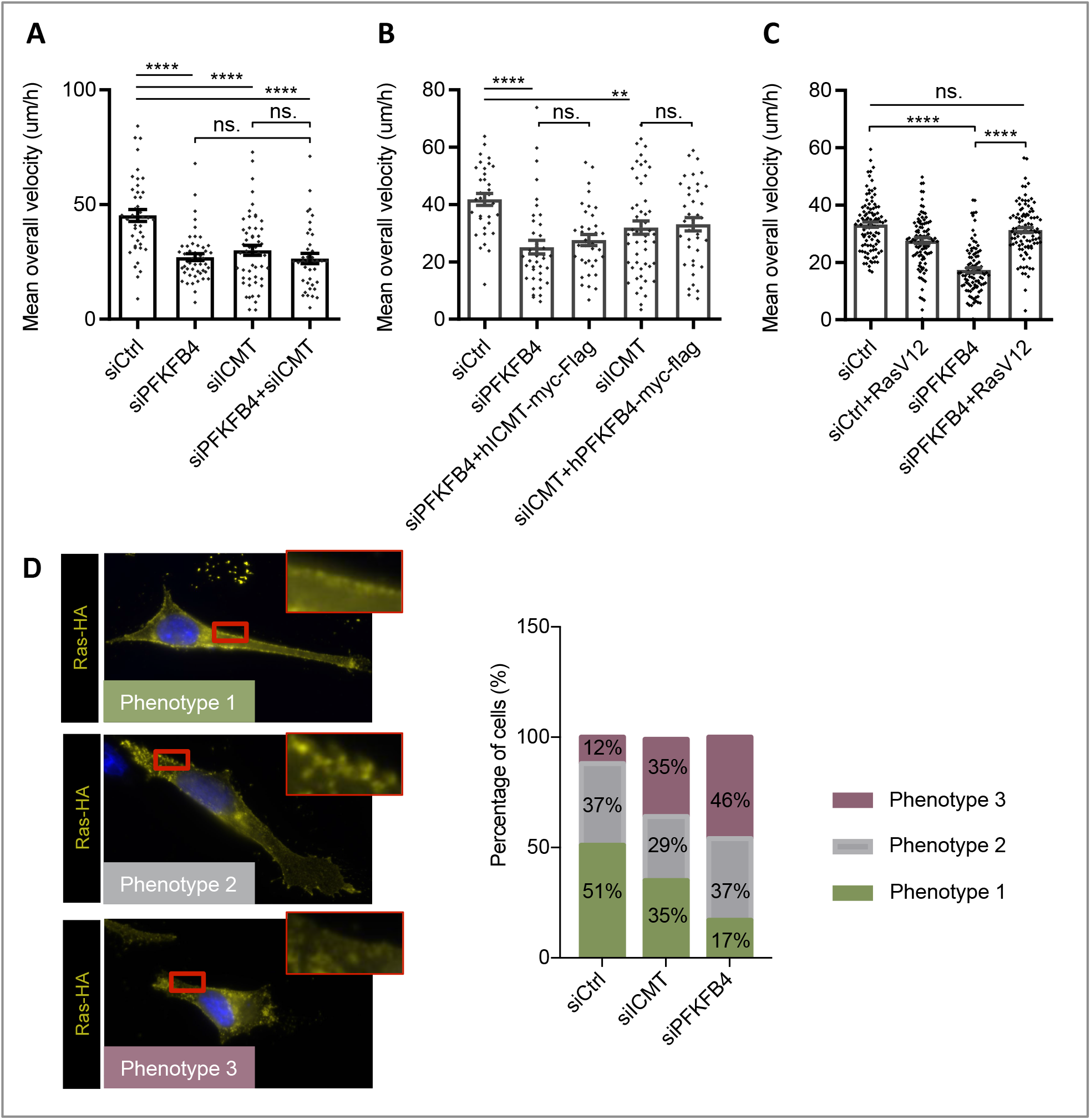
ICMT and PFKFB4 both control RAS addressing at the plasma membrane and melanoma cell migration. **(A-C)** Average speed of MeWo cells transfected with siRNA (siCtrl, siPFKFB4 or siICMT) and plasmid (empty vector, hICMT-myc-Flag, hPFKFB4-myc-Flag or RasV12). Graphs show the mean calculated in one experiment performed at least in triplicate with at least 30 cells in each condition, Error bars are calculated with SEM. p-values are calculated using the Mann-Whitney test. n.s.: p > 0,05; *: p < 0,05; **: p < 0.01; ***: p < 0.001 and ****: p < 0.0001. (**D)** Detection of RAS subcellular localization by immunostaining on MeWo cell transfected with RasV12-HA (yellow). Nuclei were stained with DAPI (blue). At the plasma membrane, RAS was found distributed according to three main phenotypes: either a clear and homogeneous membrane localization (Phenotype 1), or the absence of signal (Phenotype 3), or an intermediate phenotype with intermittent RAS expression at the membrane (Phenotype 2). Insets show enlargements of the areas framed in red. The proportion of each phenotypes was quantified after transfection of either siControl (nbcell=43), or siICMT (nbcell=51) or siPFKFB4 (nbcell=46). A representative experiment is shown, n=3.

In parallel, we compared PFKFB4 depletion phenotype to the known effect of ICMT depletion, as the major role of ICMT is to modify RAS proteins post-translationally for their efficient targeting to the plasma membrane (*43*). In order to be free from potential defects in RAS GTPase activity, we have used a constitutively active form of RAS, HA-tagged-RasV12, and tested if PFKFB4 influenced its subcellular localization, by immunofluorescence. After co-transfecting MeWo cells with HA-tagged-RasV12 and either siPFKFB4, siICMT or a control siRNA, we scored RAS localization to the plasma membrane qualitatively (Figure 3D). On top of the exogeneous RasV12 perinuclear location, the cells transfected with the control siRNA could be categorized into three groups: cells exhibiting a clear and homogeneous RAS membrane enrichment localization (phenotype 1), cells without RAS membrane enrichment (phenotype 3), and cells with an in-between phenotype with discontinuous RAS membrane localization (phenotype 2) (Figure 3D). While phenotype 2 was present in a similar proportion in each condition (siControl, siICMT or siPFKFB4; 30-37% of the cells), cells with RAS at the plasma membrane represented 51% of the siControl cells, but only 35% of siICMT cells and 17% of siPFKFB4 cells (Figure 3D). The remaining cells displayed phenotype 3. This assay thus showed that PFKFB4 depletion phenocopied ICMT loss for RAS addressing to the plasma membrane in melanoma cells.

Lastly, to test if RAS was indeed a downstream target of PFKFB4 in the control of cell migration, we measured MeWo and A375M cell migration efficiency after co-transfecting siPFKFB4 with the constitutively active RasV12. In both cell lines, RasV12 rescued the PFKFB4 migration phenotype (Figure 3C, Figure S5D). From this series of results, we concluded that the interaction between PFKFB4 and ICMT was critical for controlling RAS subcellular localisation and melanoma cell migration. PFKFB4 thus displays a novel function, important to modulate the activity of a major cell signaling pathway in cancer cells.

### PFKFB4 and ICMT control RAS-AKT signaling in melanoma

RAS signaling is a major hub for the cell to integrate multiple inputs from the extracellular cues as well as from intracellular parameters. Once activated, several downstream pathways are activated in normal as well as cancer cells. The major ones include MAP kinase, PI3 kinase and Ral signaling cascades (*48–50*). In melanoma, MAPK/Erk and PI3K/AKT pathways are frequently activated to control cell migration. Here, we next sought to further understand which of these two pathways was modulated by PFKFB4 depletion (Figure 4). While ERK phosphorylation (pERK) remained unchanged after transfection of MeWo cells with siPFKFB4 or siICMT, AKT phosphorylation was strongly decreased both on threonine 308 and on serine 473, the two major modifications leading to full activation of AKT (Figure 4A-C, Figure S5E). To extend this finding to other melanoma contexts, we examined AKT activation in three other cell lines after PFKFB4 depletion. In all cell lines, siRNA against PFKFB4 reduced target protein levels efficiently (Figure S5F,G). PhosphoAKT levels were significantly decreased in A375M and MNT1 cells while there was no significant decrease in Lu1205 (Figure 4B). This suggested that PFKFB4 influenced AKT signaling in several but not all melanoma cell contexts. Although it remains unclear why MAP kinase signaling remains unaffected while AKT pathway is decreased in this particular context, similar examples of selective activation of the PI3K-AKT pathway have been described before (*8*).

**Figure 4.**
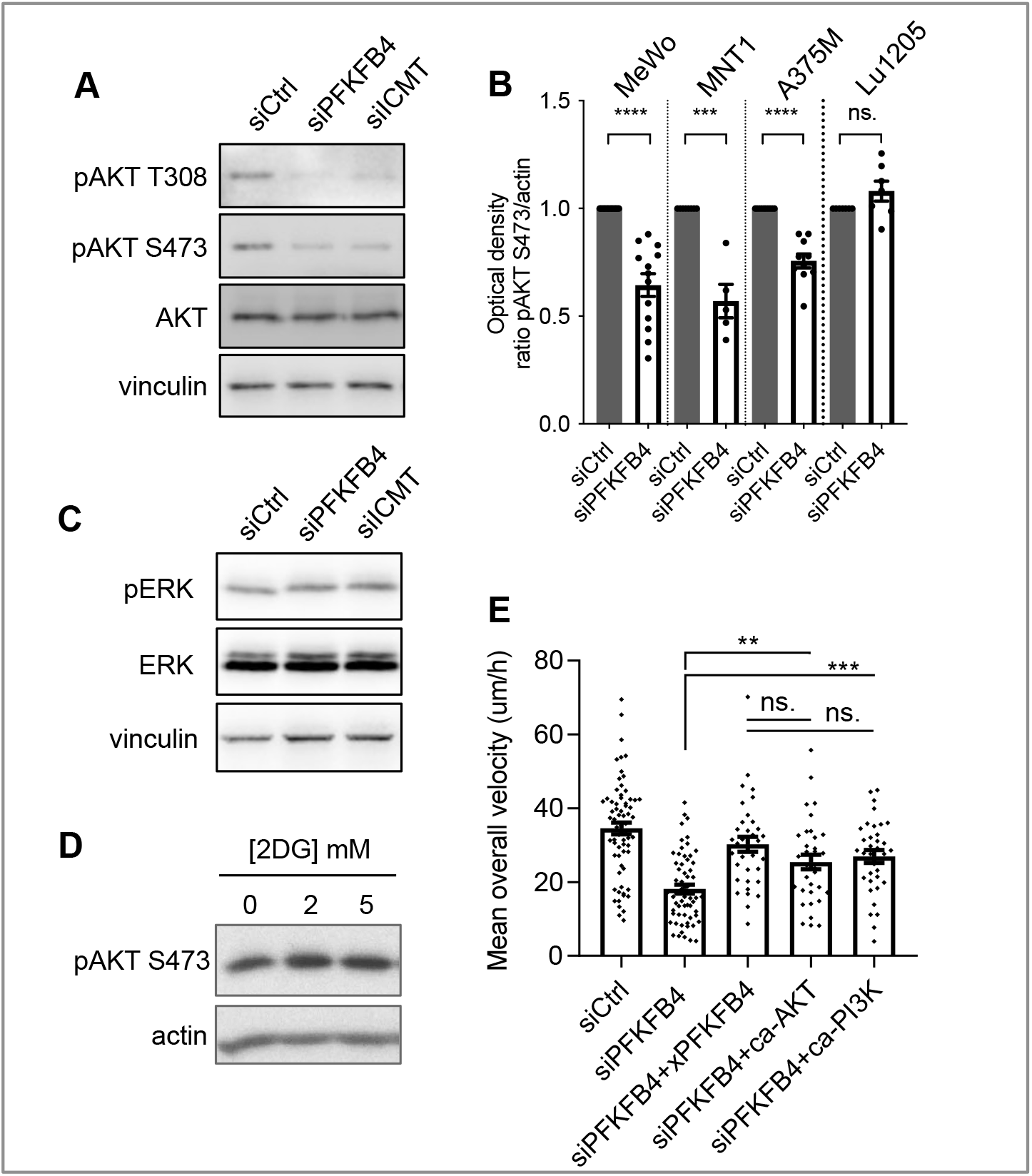
PFKFB4 and ICMT both control AKT signaling activation in melanoma cells. **(A)** Protein levels of pAKT T308, pAKT S473, AKT and actin in MeWo cells transfected with siRNA targeting PFKFB4 or ICMT. (**B**) Normalized pAKT S473/actin levels in MeWo, MNT1, A375 or Lu1205 cells transfected with siRNA targeting PFKFB4 and analyzed as in A; one point represents one biological replicate. (**C**) Protein levels of pERK, ERK and vinculin in MeWo cells transfected with siRNA targeting PFKFB4. **(D**) Protein levels of pAKT S473 in MeWo cells treated with different concentrations of 2DG for 24 hours. (**E)** Average speed of MeWo cells transfected either with siControl+empty vector, siPFKFB4+empty vector, siPFKFB4+xenopus PFKFB4 wild-type, siPFKFB4+ca-AKT, siPFKFB4+ca-PI3K. In A, C, D, E: a representative experiment is shown, n>3. Graphs show the mean ± SEM. p-values were calculated using the Mann-Whitney test. n.s. p > 0,05, ** p < 0.01, *** p < 0.001 and **** p < 0.0001.

We then wondered if PFKFB4 might affect AKT signaling as an indirect effect of glycolysis regulation. We blocked glycolysis with 2DG and tested AKT activation in MeWo cells. Although decreased lactate levels indicated an efficient block of glycolysis (Figure S3), the treatment with 2DG did not affect AKT phosphorylation on S473 (Figure 4D). As observed above for the cell migration phenotype (Figure 1), AKT phosphorylation phenotype after depleting PFKFB4 was thus not likely due to a reduction of glycolysis. Lastly, we expressed constitutively active forms of either AKT (caAKT) or its upstream regulator PI3 kinase (caPI3K) in PFKFB4-depleted MeWo cells. Defective cell migration parameters observed after PFKFB4 depletion were rescued either by caAKT or by caPI3K (Figure 4E, Figure S5H,I). All these data together indicated that PFKFB4 controlled human melanoma cell migration via a novel non-conventional function controlling the RAS/PI3K/AKT pathway.

## Discussion

PFKFB proteins are long-known major regulators of the rate of glycolysis in normal and cancer cell types (*11*). They have been involved in mediating the Warburg effect in many different tumors (*17, 19, 22, 25, 30, 51*). In particular, elevated levels of *PFKFB4* expression have been described in melanoma (Figure S1, (*30*)). Using a survey of human melanoma cell lines transcriptomes (*36*), we have selected cells with high *PFKFB4* levels, and explored PFKFB4 function in the biology of those cells, focusing on their migration in vitro. We first showed that PFKFB4 enhanced cell migration irrespective of the cells’ glycolysis levels (Figure 1, S2). Moreover, neither PFKFB4 kinase nor its phosphatase activity was required for this effect, suggesting alternative molecular mechanisms, such as protein-protein interactions. Using PFKFB4 immunoprecipitation followed by mass spectrometry, we have identified a partner of PFKFB4 which was selected for further analysis: ICMT, an enzyme embedded into the endoplasmic reticulum membrane, which adds the terminal methyl group to RAS GTPases post-translationally. This modification is required for anchorage of RAS GTPases at the plasma membrane, where RAS activates several downstream signaling events (*7*). Strikingly, both PFKFB4 and ICMT depletion displayed similar phenotypes including RAS mislocalization, decreased AKT activation and reduced cell migration (Figures 2, 3, 4). Migration phenotypes were rescued by a constitutively active form of RAS or by the constitutive activation of AKT signaling (Figure 5). In sum, our study has demonstrated a novel, glycolysis-independent function of PFKFB4, promoting the interaction between ICMT and RAS, resulting in enhanced migration of melanoma cells (Figure 5).

**Figure 5:**
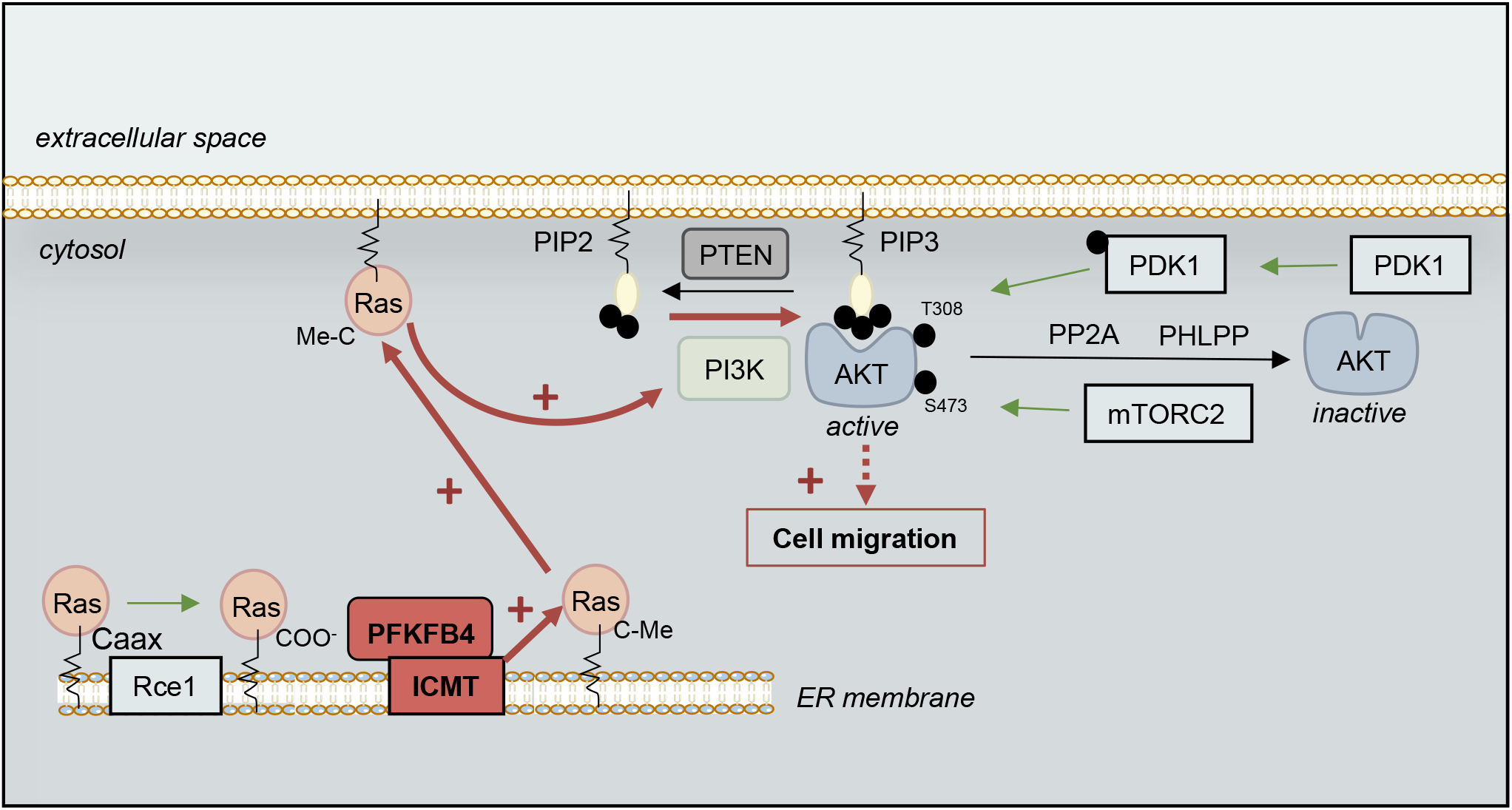
Model of cell migration control by a non-canonical function of PFKFB4, modulating RAS signaling. We propose that the interaction between PFKFB4 and ICMT would promote the ICMT/RAS interaction needed for RAS trafficking to the plasma membrane, where RAS would activate PI3K-mediated AKT phosphorylation on T308. In turn, AKT activation would then modulate cell migration. PTEN, Phosphatase and Tensin homolog; PI3K, phosphoinositide 3-kinase; PIP2, phosphatidylinositol-4,5-bisphosphate; PDK1, Phosphoinositide-dependent kinase-1; PP2A, Protein phosphatase 2; PHLPP, PH domain and Leucine rich repeat Protein Phosphatase; mTORC2, mTOR complex 2; Rce1, Ras converting enzyme 1; ICMT, isoprenylcysteine carboxyl O-methyltransferase.

Our main thought-provoking finding is that the regulation of cell migration by PFKFB4 does not depend on its kinase activity and that PFKFB4 depletion affects migration independently of experimental modulations of the high/low glycolysis status of the cells (Figures 1, S2). Our results thus uncouple the classical action of PFKFB4 in glycolysis from its role in cell migration. This undermines current strategies to counteract PFKFB4 in cancer, which involves developing pharmacological drugs to interfere with PFKFB4 kinase activity, assuming that PFKFB4’s major function in cancer is to promote glycolysis and Warburg effect (*28*). Our data show that a kinase-deficient form of PFKFB4 still retains important cancer-promoting functions outside of glycolysis regulation. Further biochemical and crystallographic analyses, beyond the scope of this cell biology study, would provide details on the protein-protein interacting subdomains interfacing PFKFB4 and ICMT. The disruption of this interaction could be an additional strategy to block PFKFB4 in cancer.

ICMT is a major posttranslational modifier of RAS GTPases. ICMT catalyzes RAS terminal methylation on the ER membrane, needed to address RAS to the plasma membrane (*41–43*). One third of human cutaneous melanoma are mutated on RAS, PI3 kinase or other partners of this pathway, enhancing its signaling activity (*5, 6*). However, here we showed that PFKFB4 promoted RAS signaling and cell migration in two cell lines with wild-type NRAS. This implied that, even in the absence of a RAS activating mutation, an increase in PFKFB4 cellular levels might also allow enhanced RAS-linked signaling and cell migration. Increase in *PFKFB4* gene expression can be achieved by hypoxia, a general feature of growing tumors. HIF1α-responsive elements have been identified to promote *pfkfb4* transcription (*18*). While in normal cells, there is a fine-tuned, dynamic and tissue-specific expression of PFKFB genes during development and cell homeostasis (*34, 52, 53*), it is likely that tumor progression enables a hypoxia-induced broader and sustained expression of PFKFB4, which would in turn promote tumor cell migration in parallel to its activation of Warburg effect. However, in melanoma cell lines, a tumor type with generally high *PFKFB4* levels, we did not observe a strict correlation between PFKFB4 expression levels and the metastatic characteristics of the cells (Figure S2). This indicates the importance of yet unknown additional cell-specific cues.

We have focused on the function of RAS-AKT signaling in the control of melanoma cell migration. This study makes a further parallel between melanoma cell features and the behavior of their parent cells in embryos, the neural crest cells. PFKFB4 was first identified as a regulator of cell migration in neural crest, and as a general patterning regulator during neural and neural crest early development (*34, 35*). Melanoma initiation and progression involves the reactivation of elements belonging to the neural crest developmental program (*54*). We here extend the parallel between the two models, showing increased expression of PFKFB4 in development and cancer as predicted by our previous WGCNA analyses (*55*). Moreover, in the embryonic cells, which rely on yolk breakdown for their energy metabolism rather than on glycolysis, we have revealed the first indications for a non-conventional function of PFKFB4. This function involved enhancing AKT signaling and cell migration (*34, 35*). It remained unclear whether this novel function of PFKFB4, found in a non-mammalian *in vivo* model, was also important in mammalian cells. Here, human melanoma cells present similar regulations by PFKFB4, implicating AKT signaling and the control of cell migration. This study thus further emphasizes the importance of PFKFB4 “moonlight” or non-conventional signaling functions, a term naming a function which is revealed when the major “sunlight” function is masked (here, the key control of glycolysis rate by PFKFB proteins).

While we stress the importance of the PFKFB4-ICMT-RAS-AKT signaling pathway, we do not exclude parallel important functions for other signaling proteins that modulate cell migration: our mass spectrometry screen for PFKFB4 partners has provided about twenty other strong interaction candidates, which could also be involved in cell trafficking or cell migration (Figures 2, S3). In conclusion, our study highlights a novel and unsuspected link between three major hallmarks of cancer cells, namely cell metabolism, signaling and migration. The crosstalk between key regulators of glycolysis and Warburg effect (PFKFB4) and a pleiotropic cell signaling pathway (ICMT-RAS) further increases the complexity of the network known to promote melanoma cancer progression. Those intricate relationships, which might also act as re-wiring options upon cancer treatment and relapse, will be important targets for future therapeutic options.

## Materials and methods

### Cloning, Plasmids

All plasmids used are listed in Table S2. For testing protein-protein interactions, a two-hybrid-like assay adapted for mammalian membrane-bound proteins (MaMTH) was used (*46, 47*). Cloning used Gibson method ((*56*), primers used in Table S3). hICMT (clone Origene n°RC207000) and hPFKFB4 (clone Origene n°RC201573) were inserted in-frame into the MaMTH bait destination vector, which contains ubiquitin C-terminal half fused to the yeast GAL4 DNA-binding domain, or into the C-tagged or N-tagged MaMTH prey destination backbone vector, which contains ubiquitin N-terminal half.

### Cell lines, cell culture, cell treatments, cell transfection

The well-characterized human melanoma cell lines MeWo (*57, 58*), A375M (*59*), MNT1 (*60*), Lu1205 (*61*) were kindly provided by Dr. L. Larue (*36*). Their mutagenic status for key driver mutations in melanoma is summarized in Figure S2 (*36, 37*). Cells were cultured in RPMI (Gibco) supplemented with 10% SVF and 1% penicillin/streptomycin (Invitrogen). HEK293T cells were cultured DMEM (Gibco) supplemented with 10% SVF and 1% penicillin/streptomycin (Invitrogen). All cell lines were incubated at 37°C with 5% CO2. At 24 hours before transfection, cell lines were plated at 200,000 cells per well (A375M and HEK293T) or 300,000 cells per well (MNT1, MeWo, Lu1205) into six-well plates. For siRNA experiments, cells were transfected either with a control siRNA (Stealth Negative Control Medium GC Duplex, Invitrogen) or with siPFKFB4 (Dharmacon Smartpool siGenome D-006764-01/02/04/17, siPFKFB4 (1); Invitrogen #HSS107863, siPFKFB4 (2)), or with siICMT (Dharmacon Smartpool siGenome #M-005209-01-0010) (Table S1) using lipofectamine RNAimax (Invitrogen) according to the manufacturer’s instructions. For the gain-of-function experiments, cells were transfected with a total of 0,5 to 1µg of DNA using lipofectamine 2000 (Invitrogen). For Check-Mate experiments, 0.5 µg of each plasmid (pBind/pG5 or pAct/pG5) were transfected using lipofectamine 2000. For glycolysis blockade, a RPMI glucose-free medium was used (Gibco). Alternatively, 2-5mM of 2-deoxy-glucose (SIGMA) was added to the normal medium. All experiments were analyzed 48h after transfection (Figures 1-4).

### Luciferase assay

A luciferase reporter driven by five GAL4 binding sites (pG5-luc) was co-transfected into HEK293T cells together with the plasmids to be tested and a control plasmid coding Renilla luciferase for normalization of the signals. Cell medium was changed 24h after transfection. At 48 hours, cells were rinsed with PBS and lysed with passive lysis buffer 1X (Promega) for 15 minutes with agitation at room temperature. Firefly and Renilla luciferase activities were measured using the Dual-Glo Luciferase Assay System (Promega). For each condition, signal intensity was normalized by the ratio between Firefly and Renilla luminescence. Transfections were performed in triplicate, each with technical duplicates.

### Protein extraction and Western-blotting

Cells were washed in PBS and lysed in RIPA buffer (10mM Tris-HCL pH 8, 150 mM NaCl, 1% NP-40, 0,1% SDS, 0,5M sodium deoxycholate) supplemented with phosphatases inhibitor (Sigma) and proteases inhibitors (Sigma) at 4°C. Protein samples were resolved on 12 % SDS-PAGE gels and transferred to PVDV membranes (Biorad). After blocking in 5% skimmed milk diluted in TBS - 0,1% Tween (TBS-T), membranes were probed with primary antibody diluted in the blocking buffer overnight at 4°C (dilutions are indicated in Table S4). After three washes in TBS-T, membranes were probed with HRP-conjugated goat anti-rabbit or anti-mouse (1:20000) 1 hour at room temperature. ECL signal was quantified by densitometric analyses using Image J software (http://rsb.info.nih.gov/ij/).

### Co-Immunoprecipitation and mass-spectrometry

At 48h after transfection, total proteins were extracted using a mild lysis buffer (100mM NaCl, 0,5% NP40, 20mM Tris-HCl pH 7,5, 5mM MgCl_2_, protease inhibitors and phosphatases inhibitors). Cells were lysed by mechanical passages through a 26-gauge syringe. For each sample, 20 µl of FLAG-M2 magnetic beads (Sigma M8823) previously washed in lysis buffer were added and incubated with agitation overnight at 4°C. Beads were then washed five times in lysis buffer. For mass spectrometry analysis, beads were further washed twice with H_2_O. Proteins were digested with trypsin, desalted using ZipTip C18 and analyzed using a nanoESI-Orbitrap Fusion (ThermoScientific). Data were analyzed using Mascot (Matrix Science). Mascot scores of the negative control, here tagged-V5 PFKFB4, were compared to the sample tagged-FLAG PFKFB4 protein, as advised by the platform. For analysis by Western-blotting, the co-immunoprecipitated proteins were eluted by boiling 10 min in Laemmli buffer (50mM Tris pH 6.8, glycerol, 2% SDS, 3% DTT, bromophenol blue), or eluted by competition with Flag peptide at 200ug/ml (five incubations of 5 minutes, P4799, Sigma). Samples were finally concentrated using 3KDa columns (Millipore).

### RNA extraction and RT-qPCR

Total RNA was extracted from cells lysed in Trizol® (Thermofisher), and then purified by chloroform extraction and isopropanol precipitation. We used M-MLV reverse transcriptase (Promega) for reverse transcription and SYBR Green mix (Biorad) for quantitative PCR. Results were normalized against reference genes *tbp* and *18S* (see Table S5 for sequences).

### L-lactate dosage

Cell supernatant was collected 24h after changing cell medium then immediately filtered by centrifugation for 15 minutes at 4°C on a 10KDa column (Abcam or Millipore). This step eliminates proteins, including Lactate Dehydrogenase (LDH) to avoid non-specific L-Lactate degradation in the sample. Extracellular L-Lactate concentration was measured from the filtered medium using L-Lactate kit (Abcam ab65330).

### Two-Dimensional random cell migration assayed by time-lapse video microscopy

We dispensed 50,000 cells per well into twelve-well plates coated with Matrigel (FisherScientific). Two-Dimensional (2D) random cell migration was monitored by time-lapse video microscopy under bright white light, with an inverted phase contrast microscope (Leica MM AF) equipped with a cell culture chamber (37°C, humidified atmosphere containing 5% CO2), an x–y–z stage controller and a charge-coupled device (CCD) CoolSnap camera (Photometrics). Images were acquired at 8-minute intervals during 16 hours, with the Metamorph software (Molecular Devices). Movies were reconstructed with the ImageJ software (http://rsbweb.nih.gov/ij/). Cells were tracked manually and parameters were calculated with ImageJ Manual Tracking plug-in.

### Immunofluorescence

MeWo cells were plated 24h before transfection on glass-coverslips. At 48h after transfection, cells were rinsed with PBS, fixed with paraformadehyde 4% for 15 minutes. After PSB wash, cells were permeabilized and non-specific protein binding blocked in 10% SVF, 0,1% Triton in PBS for 1h at room temperature. Then, cells were incubated at room temperature for 1 hour with primary antibodies diluted in blocking buffer (Table S4), rinsed with PBS, incubated 1h at room temperature in dark in secondary antibody at a 1:1000 dilution in blocking buffer (Alexa Fluor 647/555/488-conjugated goat anti rabbit/mouse/rat). The actin cytoskeleton was stained with Alexa Fluor 647 or 488 Phalloidin (Invitrogen). Cell nuclei were stained by DAPI at 1µg/ml in PBS for 10 minutes at room temperature. Coverslips were then mounted using ProLong Diamond (Molecular Probes), and imaged with 63X or 100X oil immersion objective of a widefield microscope (DM RXA, Leica; camera CoolSNAP HQ, Photometrics), using Metamorph software.

## Acknowledgements

The authors are very grateful to Drs. C. Pouponnot, M. Alkobtawi, A. Eychène, S. Saule and L. Larue for insightful scientific discussions during this study and to Dr. V. Petit for guidance in the culture of melanoma cell line. We deeply thank S. Seal, C. Pouponnot, and M. Alkobtawi for their proofreading of the manuscript. We thank all the Monsoro-Burq team members for their constant support and Drs M. Perron, P. Gilardi and C. Pouponnot for acting as thesis committee advisors. Dr. A. Eychène and Dr. L. Larue kindly provided reagents and cell lines. We thank F. Maczkowiak for subcloning the X. laevis *pfkfb4*. We also acknowledge the valuable help from M.N. Soler and L. Besse on the Institut Curie Imaging platform facility PICT-IBiSA, and members of the mass spectrometry platform of the Jacques Monod Institute (Paris). We also acknowledge the lab of Dr. Staglar who provided the constructs and cell lines for the MaMTH experiment. This work was supported by grants to A.H. M.-B. from Université Paris Saclay, Centre National de la Recherche Scientifique (CNRS), Agence Nationale pour la Recherche (ANR-15-CE13-0012-01-CRESTNETMETABO), Fondation pour la Recherche Médicale (DEQ20150331733) and Institut Universitaire de France (IUF). M.S. was supported by doctoral fellowships from Fondation Pour la Recherche Médicale (FRM ECO20160736105; FRM FDT201904007974).

## Supplementary Materials

### Supplementary Figure Legends

**Figure S1:**
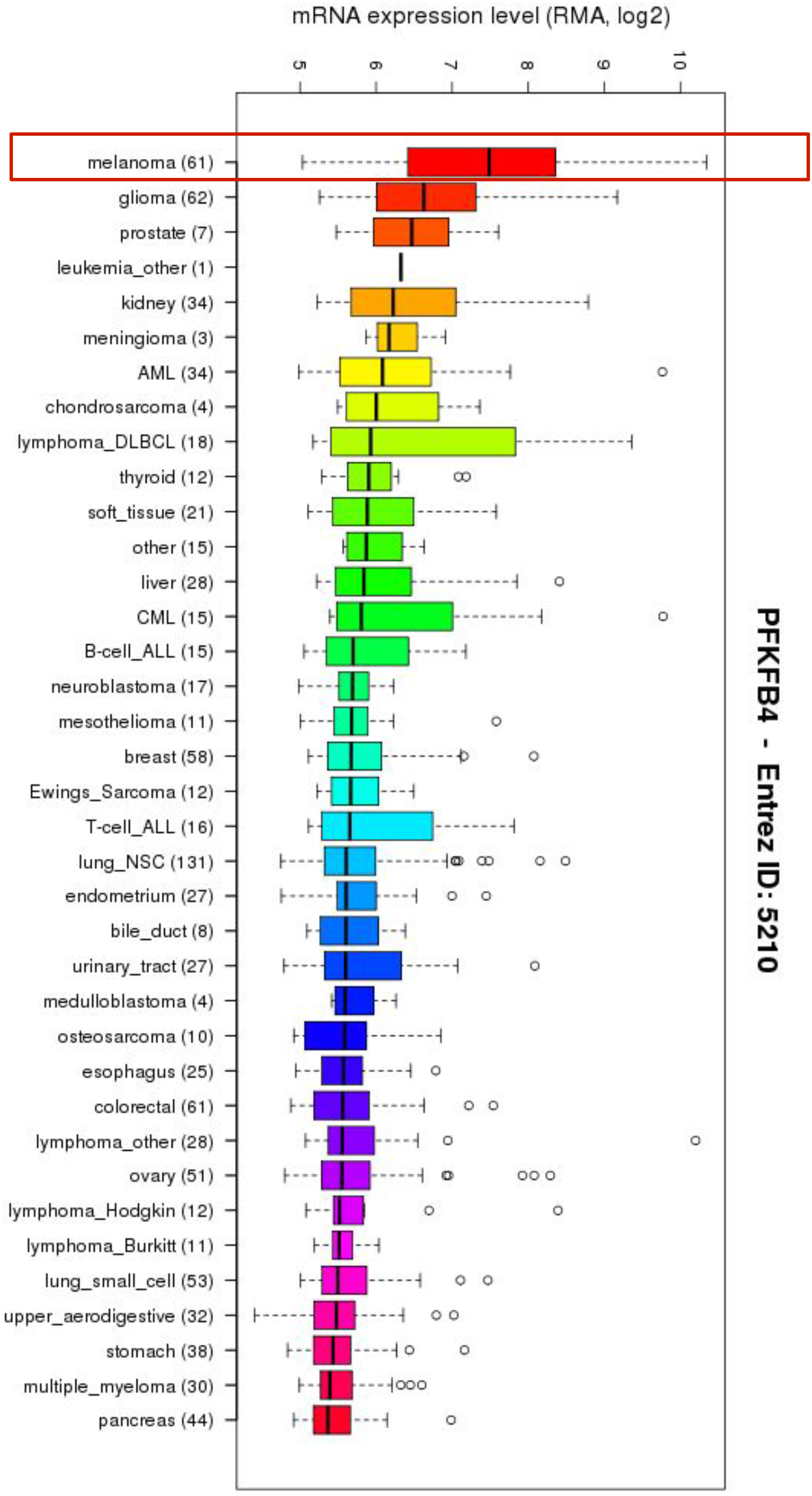
Melanomas exhibit high expression of *PFKFB4 mRNA*. Compared to other tumors, melanomas express high levels of *PFKFB4*, followed by gliomas. https://portals.broadinstitute.org/ccle/

**Figure S2:**
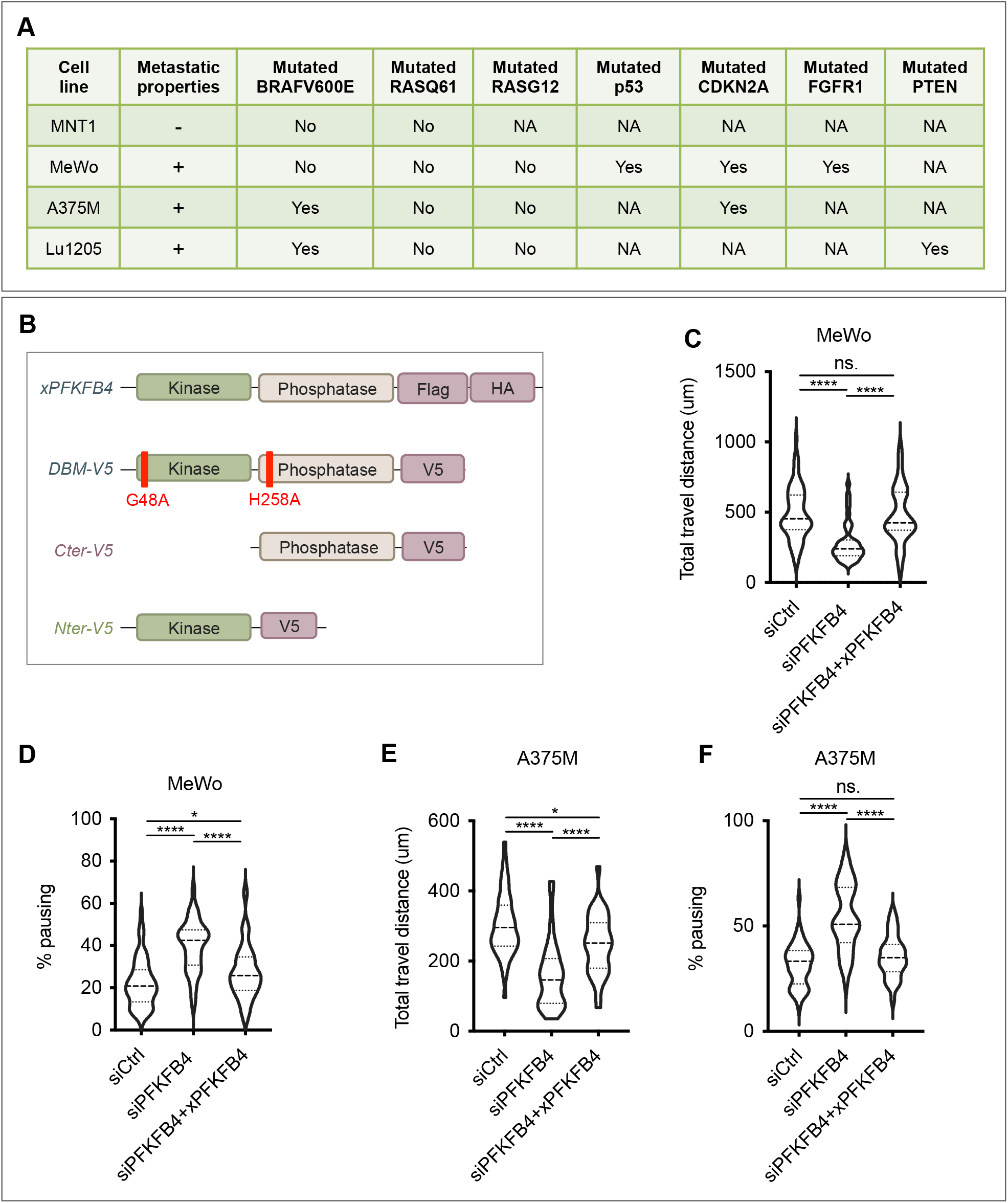
PFKFB4 controls metastatic melanoma cell migration in vitro. (A) Metastasis properties and mutational status of the cell lines used in this study. Sequencing results for key melanoma driver mutations BRafV600E and RasQ61/G12 locus were retrieved from (*36, 37*) and ATCC database. NA: non-available data. (B) Schematic structure of the *Xenopus laevis* PFKFB4 wild-type protein (xPFKFB4) and V5-tagged xPFKFB4 mutants. The *x*PFKFB4-DBM-V5 (G48A;H258A) mutant is both kinase-dead and phosphatase-dead. The xPFKFB4-Nter-V5 is a truncated form of PFKFB4 with the kinase domain conserved. xPFKFB4-Cter-V5 is a truncated form of PFKFB4 with the domain phosphatase domain conserved. Violin plots showing the travel distance (C,E) and pausing (D,F) parameters measured in MeWo (C,D) and A375M (E,F) cells, 48h after transfection either with siControl+empty vector, siPFKFB4+empty vector or siPFKFB4+xenopus PFKFB4 wild-type. The violin plots represent the probability density at each value, lines are plotted at the median and quartiles. Each graph represents one experiment performed more than three times.

**Figure S3:**
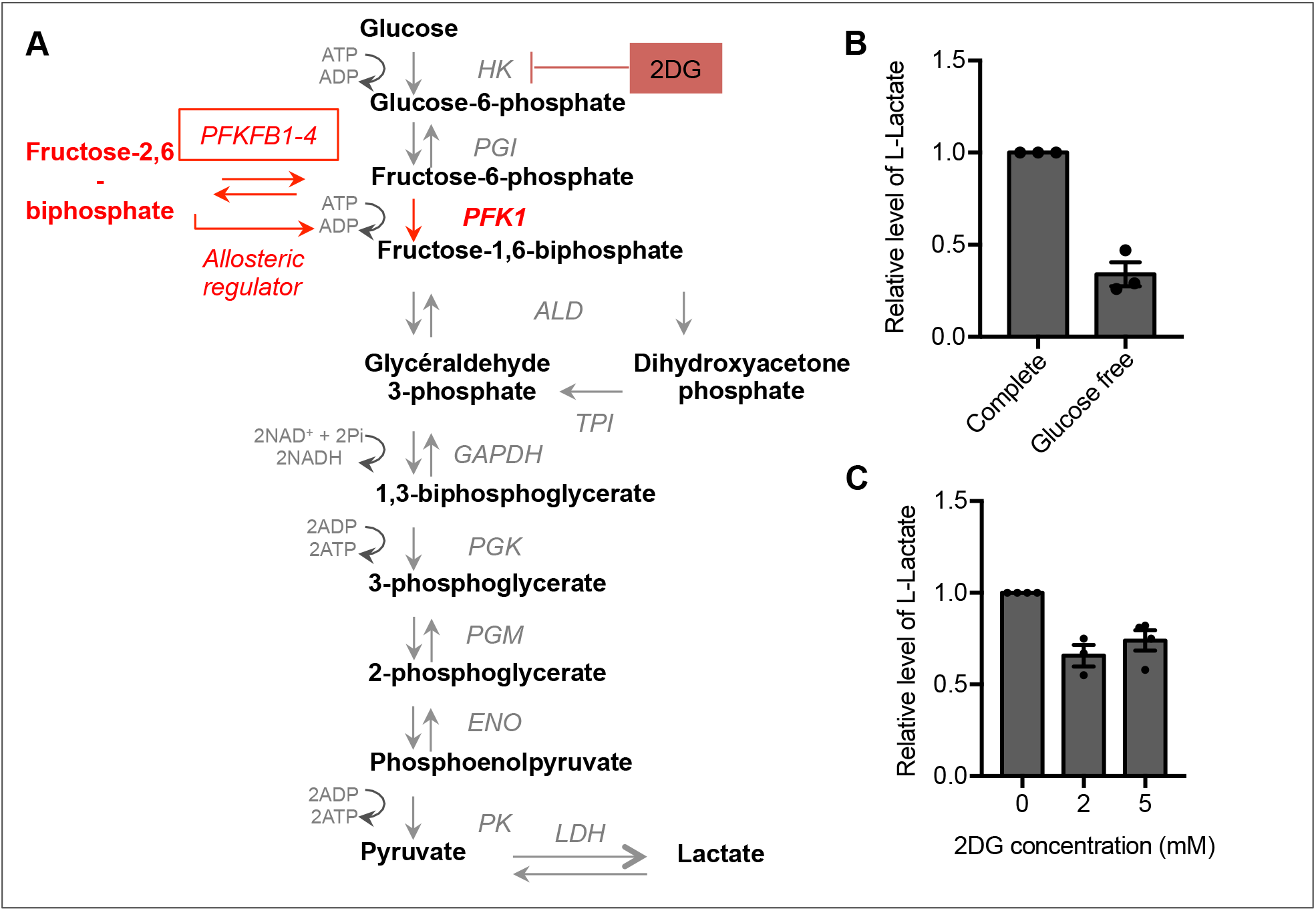
PFKFB4 controls metastatic melanoma cell migration in a glycolysis-independent manner. (A) Summary of glycolysis. The rate-limiting step of glycolysis is the second irreversible reaction, catalyzed by phosphofructokinase-1 (PFK1). PFK1 activity depends on the availability of its allosteric regulator, fructose 2,6-bisphosphate. PFKFB4 catalyses the synthesis or degradation of fructose 2,6-bisphosphate. The 2-deoxyglucose (2DG) is a glucose analog that cannot be metabolized. 2DG blocks glycolysis by competition with cellular glucose. At the end of the glycolysis, pyruvate is transformed into lactate, which is secreted in the extracellular medium. Lactate levels are a readout for the rate of glycolysis. (B) Relative extracellular L-Lactate levels in MeWo cells cultured either in a complete medium (reference) or in glucose-free medium. (C) Relative level of extracellular L-lactate of MeWo cells treated with different concentrations of 2DG.

**Figure S4:**
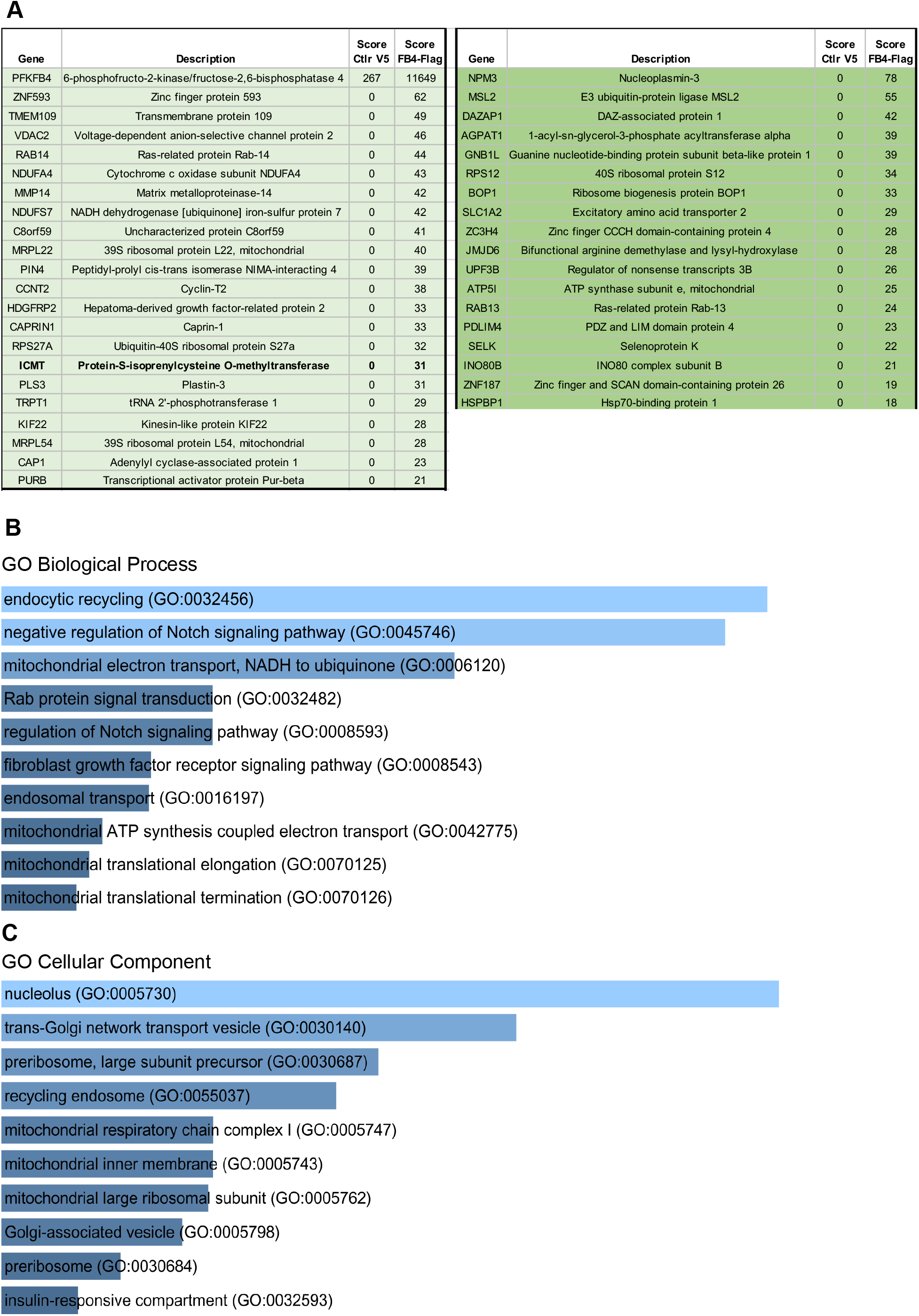
Results of IP PFKFB4 followed by mass spectrometry. (A) List of the top-41 candidate interactants of PFKFB4. We divided the result list into two parts. On the left, common targets found when the immunoprecipitation is done with xenopus or human PFKFB4 (23 targets in light green). On the right, targets only found with human PFKFB4 (18 targets in dark green). (B, C) Gene ontology analysis of potential partners of PFKFB4 – EnrichR results. (B) GO Biological process 2018 of EnrichR datas, sorted by p-value ranking. (C) GO Cellular component 2018 of EnrichR datas, sorted by p-value ranking.

**Figure S5:**
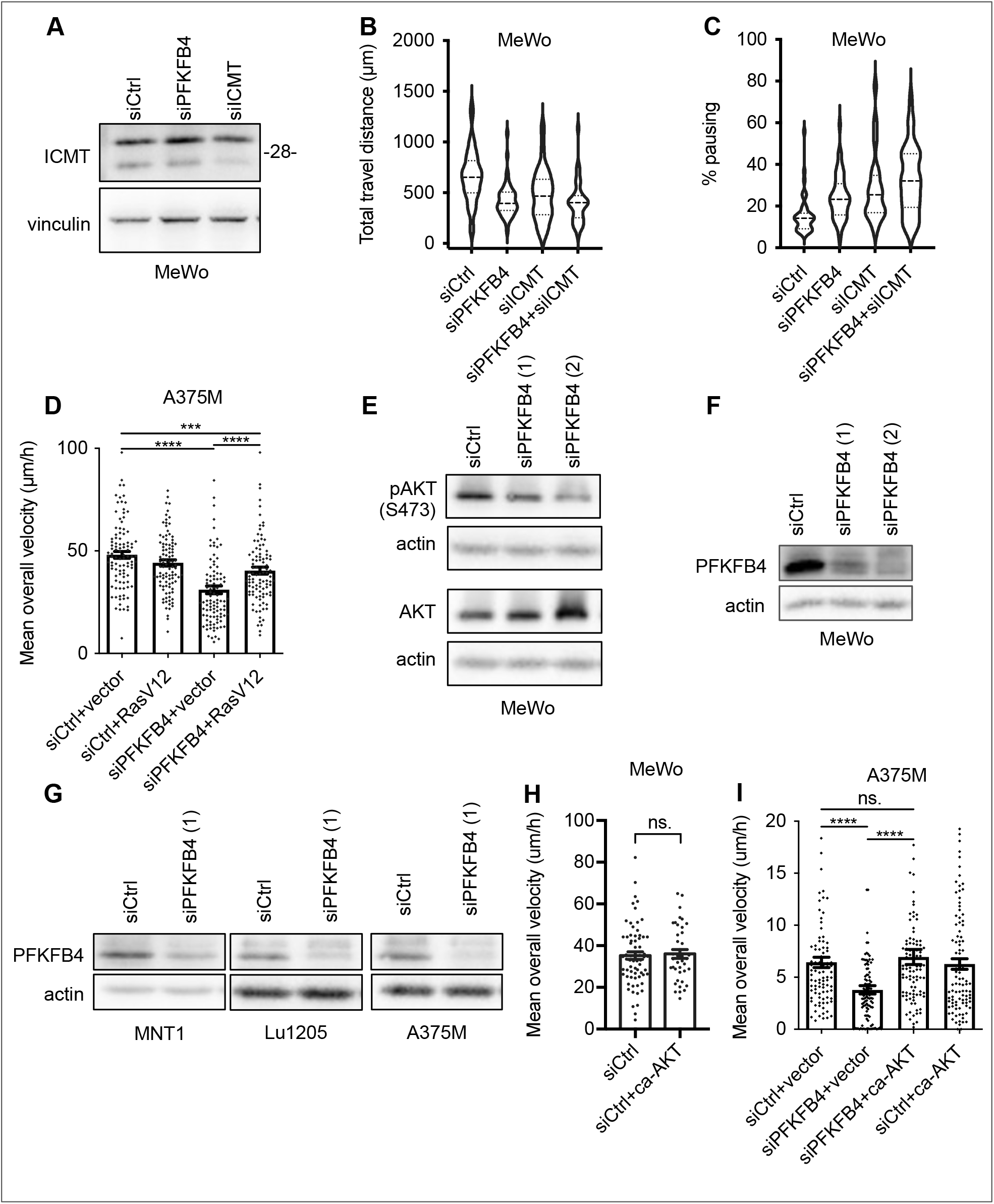
PFKFB4 and ICMT control AKT signaling in melanoma. (A) Control of the efficiency of siRNA targeting ICMT in MeWo cells. (B,C) Violin plots showing the travel distance (B) and pausing (C) parameters measured in MeWo cells, 48h after transfection either with siControl, siPFKFB4, siICMT or both. The violin plots represent the probability density at each value, lines are plotted at the median and quartiles. (D) Average speed of MeWo cells transfected with siRNA (siCtrl or siPFKFB4) and plasmid (empty vector or RasV12). (E) Protein levels pAKT S473, AKT and actin in MeWo cells transfected with two different siRNA targeting PFKFB4. (F,G) Control of the efficiency of siRNAs targeting PFKFB4 in different melanoma cell lines by Western-blot 48h after transfection. (H) Average speed of MeWo cells 48h after transfection either with siControl+empty vector or siCtrl+ca-AKT. This experiment shows that ca-AKT alone does not increase the average speed of MeWo cells. (I) Average speed of A375M cells transfected either with siControl+empty vector, siPFKFB4+empty vector, siPFKFB4+xenopus PFKFB4 wild-type, siPFKFB4+ca-AKT. In D, H, I: Graphs show the mean ± SEM. p-values were calculated using the Mann-Whitney test. n.s. p > 0,05, ** p < 0.01, *** p < 0.001 and **** p < 0.0001.

### Supplementary Tables

**Table S1:**
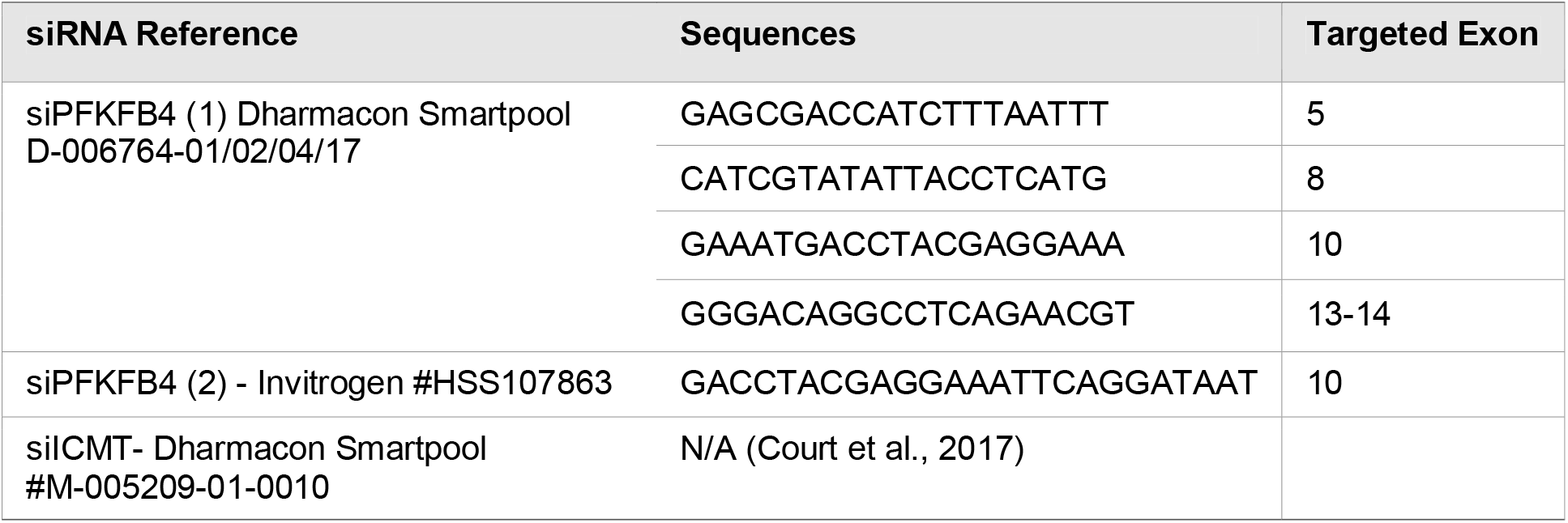
small interfering RNAs.

**Table S2:**
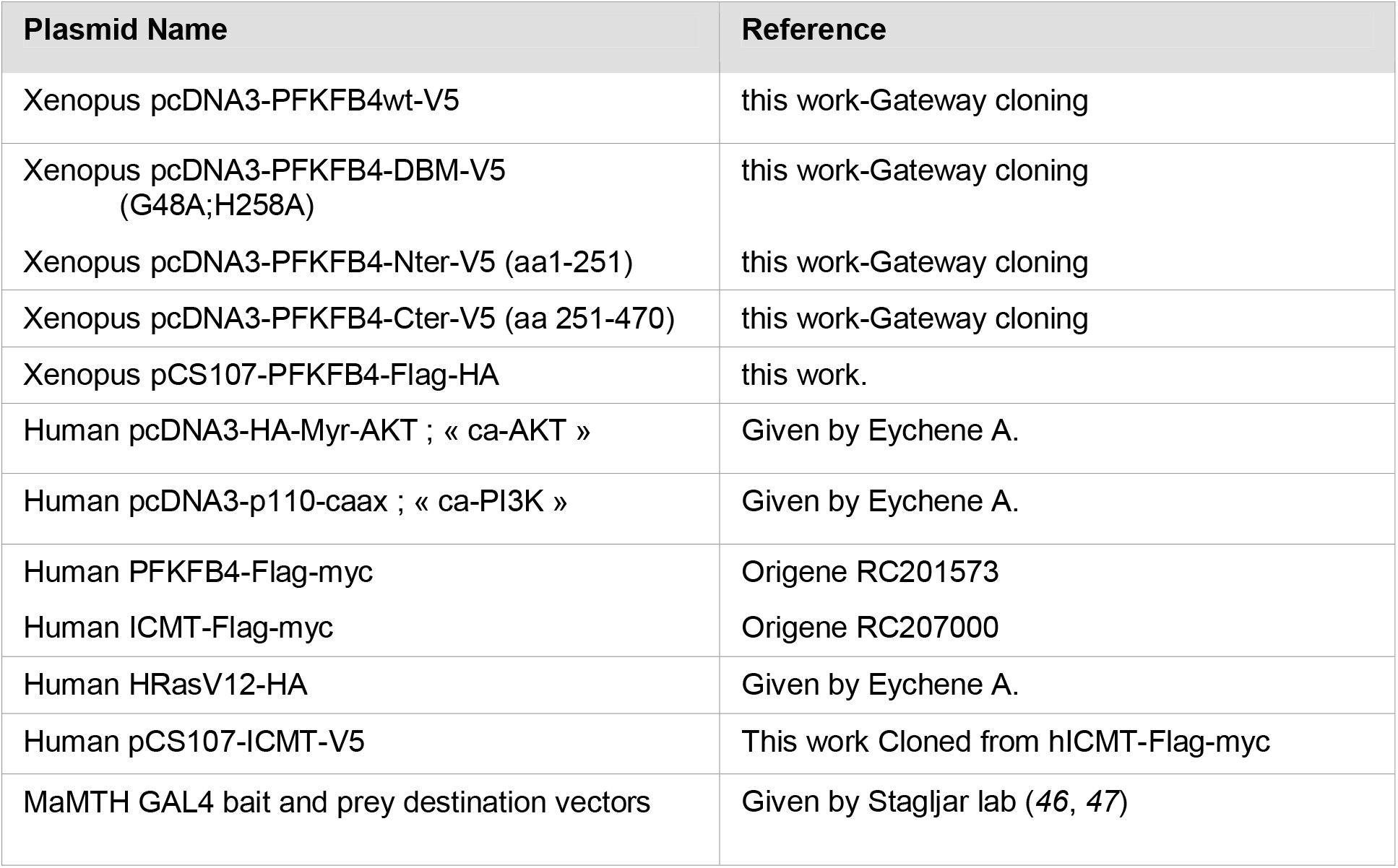
Plasmids.

**Table S3:**
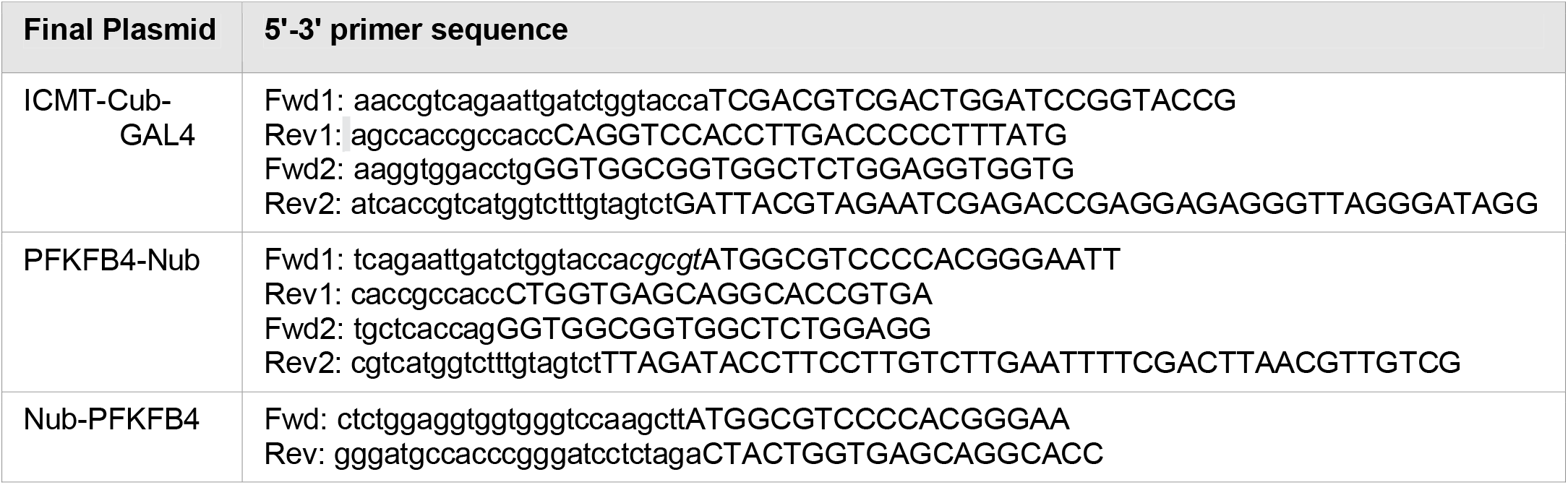
Primers used for Gibson cloning.

**Table S4:** Primary antibodies.

| Antibody name /species | Dilution | Provider | Reference |
| --- | --- | --- | --- |
| PFKFB4 Rabbit | 1/2000 | Abcam | ab137785 |
| PFKFB3 Rabbit | 1/1000 | Abcam | Ab181861 |
| Phospho-AKT (Thr308) Rabbit | 1/1000 | Cell signaling | 4056 |
| Phospho-AKT (Ser473) Rabbit | 1/1000 | Ozyme | 4060 |
| AKT Rabbit | 1/1000 | Cell signaling | 9272 |
| Actin Rabbit | 1/2000 | Sigma | A2066 |
| Vinculin HVINI Mouse | 1/5000 | Sigma | V9131 |
| Phospho-ERK 1&2 Mouse | 1/1000 | Sigma | M9692 |
| ERK Rabbit | 1/1000 | Santacruz | Sc-93 |
| Phospho-PDK1 (Ser241) Rabbit | 1/1000 | Cell signaling | 3061 |
| PDK1 Rabbit | 1/1000 | Cell signaling | 3062 |
| PTEN Rabbit | 1/1000 | Cell signaling | 9559 |
| Phospho-GSK-3beta (Ser9) Rabbit | 1/1000 | Ozyme | 9336 |
| ICMT Rabbit | 1/1000 | Proteintech | 51001-2-AP |
| HA Rat | 1/1000 for WB;<br>1/500 for IF | Roche | Clone 3F10 |
| Flag Rabbit | 1/1000 | Sigma | F7425 |

**Table S5:**
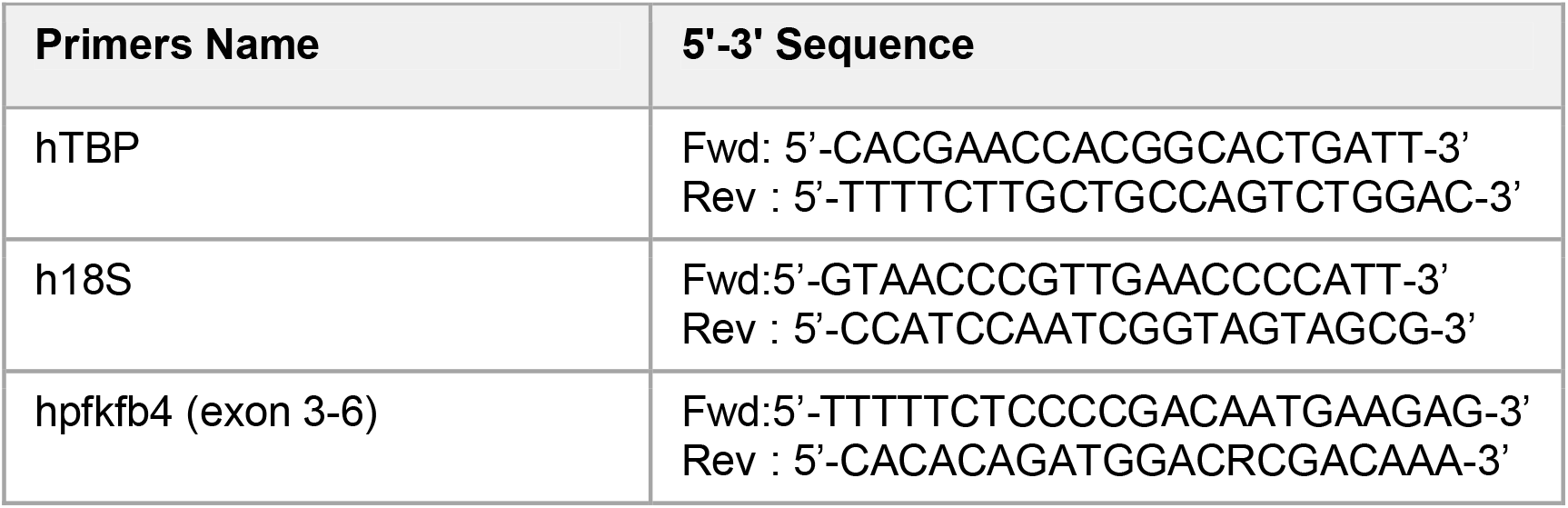
PCR primers.

## References

1. E. T. Roussos, J. S. Condeelis, A. Patsialou, Chemotaxis in cancer. Nat. Rev. Cancer. 11, 573–587 (2011).

2. E. Theveneau, R. Mayor, in Neural Crest Cells (Elsevier, 2014; http://linkinghub.elsevier.com/retrieve/pii/B9780124017306000041), xpp. 73–88.

3. D. Hanahan, R. A. Weinberg, Hallmarks of cancer: The next generation. Cell. 144, 646–674 (2011).

4. C. Bertolotto, Melanoma: From Melanocyte to Genetic Alterations and Clinical Options. Scientifica (Cairo). 2013, 1–22 (2013).

5. .The Cancer Genome Atlas Network, Genomic Classification of Cutaneous Melanoma. Cell. 161, 1681–1696 (2015).

6. Z. Ali, N. Yousaf, J. Larkin, Melanoma epidemiology, biology and prognosis. Eur. J. Cancer, Suppl. 11, 81–91 (2013).

7. A. D. Cox, C. J. Der, Ras history. Small GTPases. 1, 2–27 (2010).

8. C. Posch, M. Sanlorenzo, I. Vujic, J. A. Oses-Prieto, B. D. Cholewa, S. T. Kim, J. Ma, K. Lai, M. Zekhtser, R. Esteve-Puig, G. Green, S. Chand, A. L. Burlingame, R. Panzer-Grümayer, K. Rappersberger, S. Ortiz-Urda, Phosphoproteomic Analyses of NRAS(G12) and NRAS(Q61) Mutant Melanocytes Reveal Increased CK2α Kinase Levels in NRAS(Q61) Mutant Cells. J. Invest. Dermatol. 136, 2041–2048 (2016).

9. M. V Liberti, J. W. Locasale, The Warburg Effect: How Does it Benefit Cancer Cells? Trends Biochem. Sci. 41, 211–218 (2016).

10. H. G. Hers, E. Van Schaftingen, Fructose 2,6-bisphosphate 2 years after its discovery. Biochem. J. 206, 1–12 (1982).

11. M. H. Rider, L. Bertrand, D. Vertommen, P. A. Michels, G. G. Rousseau, L. Hue, 6-Phosphofructo-2-Kinase/Fructose-2,6-Bisphosphatase: Head-To-Head With a Bifunctional Enzyme That Controls Glycolysis. Biochem. J. 381, 561–79 (2004).

12. P. Bruni, P. Vandoolaeghe, G. G. Rousseau, L. Hue, M. H. Rider, Expression and regulation of 6-phosphofructo-2-kinase/fructose-2,6-bisphosphatase isozymes in white adipose tissue. Eur. J. Biochem. 259, 756–61 (1999).

13. N. P. Manes, M. R. El-Maghrabi, The kinase activity of human brain 6-phosphofructo-2-kinase/fructose-2,6-bisphosphatase is regulated via inhibition by phosphoenolpyruvate. Arch. Biochem. Biophys. 438, 125–136 (2005).

14. S. J. Pilkis, T. H. Claus, I. J. Kurland, A. J. Lange, 6-Phosphofructo-2-Kinase/Fructose-2,6-Bisphosphatase: A Metabolic Signaling Enzyme. Annu. Rev. Biochem. 64, 799–835 (1995).

15. M. H. Rider, J. Vandamme, E. Lebeau, D. Vertommen, H. Vidal, G. G. Rousseau, J. Vandekerckhove, L. Hue, The two forms of bovine heart 6-phosphofructo-2-kinase/fructose-2,6-bisphosphatase result from alternative splicing. Biochem. J. 285 (Pt 2, 405–11 (1992).

16. E. van Schaftingen, B. Lederer, R. Bartrons, H.-G. Hers, A kinetic study of pyrophosphate: fructose 6-phosphate phosphotransferase from potato tubers. Eur.J.Biochem. 129, 191–195 (1982).

17. O. H. Minchenko, A. Ochiai, I. L. Opentanova, T. Ogura, D. O. Minchenko, J. Caro, S. V. Komisarenko, H. Esumi, Overexpression of 6-phosphofructo-2-kinase/fructose-2,6-bisphosphatase-4 in the human breast and colon malignant tumors. Biochimie. 87, 1005–1010 (2005).

18. O. H. Minchenko, I. Opentanova, D. Minchenko, T. Ogura, H. Esumi, Hypoxia induces transcription of 6-phosphofructo-2-kinase/fructose-2,6-biphosphatase-4 gene via hypoxia-inducible factor-1α activation. FEBS Lett. 576, 14–20 (2004).

19. O. H. Minchenko, K. Tsuchihara, D. O. Minchenko, A. Bikfalvi, H. Esumi, Mechanisms of regulation of PFKFB expression in pancreatic and gastric cancer cells. World J. Gastroenterol. 20, 13705–13717 (2014).

20. V. Goidts, J. Bageritz, L. Puccio, S. Nakata, M. Zapatka, S. Barbus, G. Toedt, B. Campos, A. Korshunov, S. Momma, E. Van Schaftingen, G. Reifenberger, C. Herold-Mende, P. Lichter, B. Radlwimmer, RNAi screening in glioma stem-like cells identifies PFKFB4 as a key molecule important for cancer cell survival. Oncogene. 31, 3235–3243 (2012).

21. A. Houddane, L. Bultot, L. Novellasdemunt, M. Johanns, M.-A. Gueuning, D. Vertommen, P. G. Coulie, R. Bartrons, L. Hue, M. H. Rider, Role of Akt/PKB and PFKFB isoenzymes in the control of glycolysis, cell proliferation and protein synthesis in mitogen-stimulated thymocytes. Cell. Signal. 34, 23–37 (2017).

22. J. Chesney, J. Clark, A. C. Klarer, Y. Imbert-Fernandez, A. N. Lane, S. Telang, Fructose-2,6-Bisphosphate synthesis by 6-Phosphofructo-2-Kinase/Fructose-2,6-Bisphosphatase 4 (PFKFB4) is required for the glycolytic response to hypoxia and tumor growth. Oncotarget. 5, 6670–6686 (2014).

23. X. Zhang, X. Yang, C. Yang, P. Li, W. Yuan, Targeting protein kinase CK2 suppresses bladder cancer cell survival via the glucose metabolic pathway. Oncotarget (2016).

24. S. Ros, C. R. Santos, S. Moco, F. Baenke, G. Kelly, M. Howell, N. Zamboni, A. Schulze, Functional metabolic screen identifies 6-phosphofructo-2-kinase/fructose-2, 6-biphosphatase 4 as an important regulator of prostate cancer cell survival. Cancer Discov. 2, 328–343 (2012).

25. S. J. Yun, S. W. Jo, Y. S. Ha, O. J. Lee, W. T. Kim, Y. J. Kim, S. C. Lee, W. J. Kim, PFKFB4 as a prognostic marker in non-muscle-invasive bladder cancer. Urol. Oncol. Semin. Orig. Investig. 30, 893–899 (2012).

26. Y. Shu, Y. Lu, X. Pang, W. Zheng, Y. Huang, J. Li, J. Ji, C. Zhang, P. Shen, Phosphorylation of PPARγ at Ser84 promotes glycolysis and cell proliferation in hepatocellular carcinoma by targeting PFKFB4. Oncotarget. 7 (2016), doi:10.18632/oncotarget.12764.

27. L. Yao, L. Wang, Z.-G. Cao, X. Hu, Z.-M. Shao, High expression of metabolic enzyme PFKFB4 is associated with poor prognosis of operable breast cancer. Cancer Cell Int. 19, 165 (2019).

28. J. Chesney, J. Clark, L. Lanceta, J. O. Trent, S. Telang, Targeting the sugar metabolism of tumors with a first-in-class 6-phosphofructo-2-kinase (PFKFB4) inhibitor. Oncotarget. 6 (2015), doi:10.18632/oncotarget.4534.

29. A. Houddane, L. Bultot, L. Novellasdemunt, M. Johanns, M.-A. Gueuning, D. Vertommen, P. G. Coulie, R. Bartrons, L. Hue, M. H. Rider, Role of Akt/PKB and PFKFB isoenzymes in the control of glycolysis, cell proliferation and protein synthesis in mitogen-stimulated thymocytes. Cell. Signal. 34, 23–37 (2017).

30. O. H. Minchenko, T. Ogura, I. L. Opentanova, D. O. Minchenko, H. Esumi, Splice isoform of 6-phosphofructo-2-kinase/ fructose-2,6-bisphosphatase-4: Expression and hypoxic regulation. Mol. Cell. Biochem. 280, 227–234 (2005).

31. Q. Wang, F. Zeng, Y. Sun, Q. Qiu, J. Zhang, W. Huang, J. Huang, X. Huang, L. Guo, Etk interaction with PFKFB4 modulates chemoresistance of small-cell lung cancer by regulating autophagy. Clin. Cancer Res. 24, 950–962 (2018).

32. A. M. Strohecker, S. Joshi, R. Possemato, R. T. Abraham, D. M. Sabatini, E. White, Identification of 6-phosphofructo-2-kinase/fructose-2,6-bisphosphatase as a novel autophagy regulator by high content shRNA screening. Oncogene. 34, 5662–5676 (2015).

33. S. Dasgupta, K. Rajapakshe, B. Zhu, B. C. Nikolai, P. Yi, N. Putluri, J. M. Choi, S. Y. Jung, C. Coarfa, T. F. Westbrook, X. H.-F. Zhang, C. E. Foulds, S. Y. Tsai, M.-J. Tsai, B. W. O’Malley, Metabolic enzyme PFKFB4 activates transcriptional coactivator SRC-3 to drive breast cancer. Nature. 4 (2018), doi:10.1038/s41586-018-0018-1.

34. C. Pegoraro, A. L. Figueiredo, F. Maczkowiak, C. Pouponnot, A. Eychene, A. H. Monsoro-Burq, PFKFB4 controls embryonic patterning via Akt signalling independently of glycolysis. Nat. Commun. 6, 5953 (2015).

35. A. L. Figueiredo, F. Maczkowiak, C. Borday, P. Pla, M. Sittewelle, C. Pegoraro, A. H. Monsoro-Burq, PFKFB4 control of AKT signaling is essential for premigratory and migratory neural crest formation. Development. 144, 4183–4194 (2017).

36. F. Rambow, B. Job, V. Petit, F. Gesbert, V. Delmas, H. Seberg, G. Meurice, E. Van Otterloo, P. Dessen, C. Robert, D. Gautheret, R. A. Cornell, A. Sarasin, L. Larue, New Functional Signatures for Understanding Melanoma Biology from Tumor Cell Lineage-Specific Analysis. Cell Rep. 13, 840–853 (2015).

37. M. Ranzani, C. Alifrangis, D. Perna, K. Dutton-Regester, A. Pritchard, K. Wong, M. Rashid, C. D. Robles-Espinoza, N. K. Hayward, U. Mcdermott, M. Garnett, D. J. Adams, BRAF/NRAS wild-type melanoma, NF1 status and sensitivity to trametinib. Pigment Cell Melanoma Res. 28, 117–119 (2015).

38. D. A. Okar, A. J. Lange, À. Manzano, A. Navarro-Sabatè, L. Riera, R. Bartrons, PFK-2/FBPase-2: Maker and breaker of the essential biofactor fructose-2,6-bisphosphate. Trends Biochem. Sci. 26 (2001), pp. 30–35.

39. L. Li, K. Lin, I. J. Kurland, J. J. Correiap, S. J. Pilkis, Site-directed Mutagenesis in Rat Liver 6-Phosphofructo-2-kinase. J. Biol. Chem. 267, 4386–4393 (1992).

40. A. Tauler, K. Lin, J. Pilkis, Hepatic 6-Phosphofructo-2-kinase/Fructose-2,6-bisphosphatase. J. Biol. Chem. 265, 15617–15622 (1990).

41. E. Choy, V. K. Chiu, J. Silletti, M. Feoktistov, T. Morimoto, D. Michaelson, I. E. Ivanov, M. R. Philips, Endomembrane trafficking of ras: The CAAX motif targets proteins to the ER and Golgi. Cell. 98, 69–80 (1999).

42. Q. Dai, E. Choy, V. Chiu, J. Romano, S. R. Slivka, S. A. Steitz, S. Michaelis, M. R. Philips, Mammalian Prenylcysteine Carboxyl Methyltransferase Is in the Endoplasmic Reticulum. J. Biol. Chem. 273, 15030–15034 (1998).

43. D. Michaelson, W. Ali, V. K. Chiu, M. Bergo, J. Silletti, L. Wright, S. G. Young, M. Philips, Postprenylation CAAX Processing Is Required for Proper Localization of Ras but Not Rho GTPases. Mol. Biol. Cell. 16, 1606–1616 (2005).

44. L. P. Wright, H. Court, A. Mor, I. M. Ahearn, P. J. Casey, M. R. Philips, Topology of Mammalian Isoprenylcysteine Carboxyl Methyltransferase Determined in Live Cells with a Fluorescent Probe. Mol. Cell. Biol. 29, 1826–1833 (2009).

45. J. Cansado, To finish things well: cysteine methylation ensures selective GTPase membrane localization and signalling. Curr. Genet. 64, 341–344 (2018).

46. P. Saraon, I. Grozavu, S. H. Lim, J. Snider, Z. Yao, I. Stagljar, Detecting Membrane Protein-protein Interactions Using the Mammalian Membrane Two-hybrid (MaMTH) Assay. Curr. Protoc. Chem. Biol. 9, 38–54 (2017).

47. J. Petschnigg, B. Groisman, M. Kotlyar, M. Taipale, Y. Zheng, C. F. Kurat, A. Sayad, J. R. Sierra, M. M. Usaj, J. Snider, A. Nachman, I. Krykbaeva, M. S. Tsao, J. Moffat, T. Pawson, S. Lindquist, I. Jurisica, I. Stagljar, The mammalian-membrane two-hybrid assay (MaMTH) for probing membrane-protein interactions in human cells. Nat. Methods. 11, 585–592 (2014).

48. P. Rodriguez-Viciana, P. H. Warne, R. Dhand, B. Vanhaesebroeck, I. Gout, M. J. Fry, M. D. Waterfield, J. Downward, Phosphatidylinositol-3-OH kinase direct target of Ras. Nature. 370 (1994), pp. 527–532.

49. T. Urano, R. Emkey, L. A. Feig, Ral-GTPases mediate a distinct downstream signaling pathway from Ras that facilitates cellular transformation. EMBO J. 15, 810–816 (1996).

50. C. Peyssonnaux, S. Provot, M. P. Felder-Schmittbuhl, G. Calothy, A. Eychène, Induction of postmitotic neuroretina cell proliferation by distinct Ras downstream signaling pathways. Mol. Cell. Biol. 20, 7068–79 (2000).

51. H. Zhang, C. Lu, M. Fang, W. Yan, M. Chen, Y. Ji, S. He, T. Liu, T. Chen, J. Xiao, HIF-1α activates hypoxia-induced PFKFB4 expression in human bladder cancer cells. Biochem. Biophys. Res. Commun. 476, 146–152 (2016).

52. C. Pegoraro, F. Maczkowiak, A. H. Monsoro-Burq, Pfkfb (6-phosphofructo-2-kinase/fructose-2,6-bisphosphatase) isoforms display a tissue-specific and dynamic expression during Xenopus laevis development. Gene Expr. Patterns. 13, 203–211 (2013).

53. O. Minchenko, I. Opentanova, J. Caro, Hypoxic regulation of the 6-phosphofructo-2-kinase/fructose-2,6-bisphosphatase gene family (PFKFB-1-4) expression in vivo. FEBS Lett. 554, 264–270 (2003).

54. C. K. Kaufman, C. Mosimann, Z. P. Fan, S. Yang, A. J. Thomas, J. Ablain, J. L. Tan, R. D. Fogley, E. van Rooijen, E. J. Hagedorn, C. Ciarlo, R. M. White, D. A. Matos, A. C. Puller, C. Santoriello, E. C. Liao, R. A. Young, L. I. Zon, E. Van Rooijen, C. Ciarlo, R. M. White, D. A. Matos, A. C. Puller, A zebrafish melanoma model reveals emergence of neural crest identity during melanoma initiation. Science (80-.). 351, aad2197 (2016).

55. J.-L. Plouhinec, S. Medina-Ruiz, C. Borday, E. Bernard, J.-P. Vert, M. B. Eisen, R. M. Harland, A. H. Monsoro-Burq, A molecular atlas of the developing ectoderm defines neural, neural crest, placode, and nonneural progenitor identity in vertebrates. PLOS Biol. 15, e2004045 (2017).

56. H. O. Smith, R.-Y. Chuang, D. G. Gibson, L. Young, C. A. Hutchison, J. C. Venter, Enzymatic assembly of DNA molecules up to several hundred kilobases. Nat. Methods. 6, 343–345 (2009).

57. M. Ishikawa, J. Dennis, S. Man, R. S. Kerbel, Isolation and Characterization of Spontaneous Wheat Germ Agglutinin-resistant Human Melanoma Mutants Displaying Remarkably Different Metastatic Profiles in Nude Mice. Cancer Res. 48, 584–588 (1988).

58. R. S. Kerbel, M. S. Man, D. Dexter, A model of human cancer metastasis: Extensive spontaneous and artificial metastasis of a human pigmented melanoma and derived variant sublines in nude mice. J. Natl. Cancer Inst. 72, 93–108 (1984).

59. P. Sriramarao, M. A. Bourdon, Melanoma cell invasive and metastatic potential correlates with endothelial cell reorganization and tenascin expression. Endothel. J. Endothel. Cell Res. 4, 85–97 (1996).

60. M. Cuomo, M. R. Nicotra, C. Apollonj, R. Fraioli, P. Giacomini, P. G. Natali, Production and Characterization of the Murine Monoclonal Antibody 2G10 to a Human T4-Tyrosinase Epitope. J. Invest. Dermatol. 96, 446–451 (1991).

61. I. Juhasz, S. M. Albelda, D. E. Elder, G. F. Murphy, K. Adachi, D. Herlyn, I. T. Valyi-Nagy, M. Herlyn, Growth and invasion of human melanomas in human skin grafted to immunodeficient mice. Am. J. Pathol. 143, 528–537 (1993).

